# The prevalence and impact of transient species in ecological communities

**DOI:** 10.1101/163816

**Authors:** Sara Snell, Brian S. Evans, Ethan P. White, Allen H. Hurlbert

## Abstract

Transient species occur infrequently in a community over time and do not maintain viable local populations. Because transient species interact differently than non-transients with their biotic and abiotic environment, it is important to characterize the prevalence of these species and how they impact our understanding of ecological systems. We quantified the prevalence and impact of transient species in communities using data on over 17,000 community time series spanning an array of ecosystems, taxonomic groups, and spatial scales. We found that transient species are a general feature of communities regardless of taxa or ecosystem. The proportion of these species decreases with spatial scale leading to a need to control for scale in comparative work. Removing transient species from analyses influences the form of a suite of commonly studied ecological patterns including species-abundance distributions, species-energy relationships, species-area relationships, and temporal turnover. Careful consideration should be given to whether transient species are included in analyses depending on the theoretical and practical relevance of these species for the question being studied.

## Introduction

Ecologists frequently conduct taxonomic surveys to characterize the diversity and composition of ecological assemblages. While many of the species observed in these surveys represent local populations, some may be irregular visitors that do not maintain viable local populations, are poorly suited to the local conditions, and rarely interact with other members of the community. Grinnell (1922) first coined the term “accidental” to refer to this kind of species, which is observed inconsistently at a site over time in contrast to the more regular and predictable members of an assemblage. This group of species has also been referred to as “occasional”, “vagrant”, “transient”, and “tourist” (Southwood et al. 1982; Costello and Myers 1996; Novotný and Basset 2000; Magurran and Henderson 2003; Ulrich and Ollik 2004; Dolan et al. 2009; Coyle et al. 2013; Petersen et al. 2015; Supp et al. 2015). Regardless of the name applied, these species (hereafter “transients”) have generally been identified based on the low frequency of observations recorded in samples or surveys over time at a given location (i.e., low temporal occupancy). Several studies ranging from birds to fish to amphipods have found that temporal occupancy is frequently bimodally distributed within communities, with one distinct mode at very low occupancy reflecting transient species, and another mode at high occupancy reflecting temporally persistent “core” species (Figure 1A; Costello and Myers 1996; Magurran and Henderson 2003; Coyle et al. 2013; Umaña et al. 2017). The existence of a mode at low occupancy indicates that transient species may make up a larger proportion of ecological assemblages than has typically been acknowledged.

**Figure 1.**
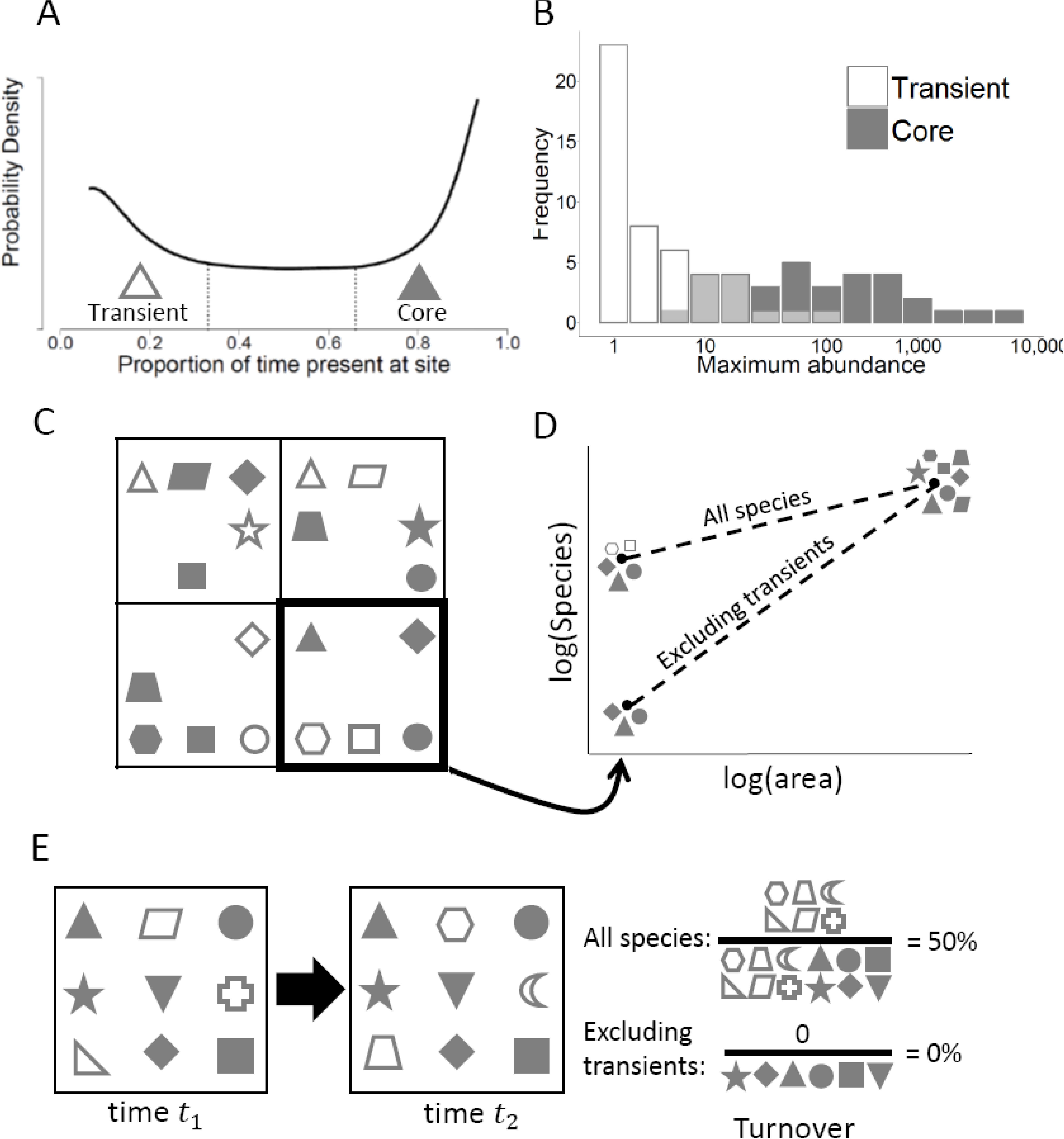
(A) Bimodal distribution of temporal occupancy for North American birds from Coyle et al. (2013) illustrating one mode of “core” species observed consistently at sites and a mode of low occupancy “transient” species observed irregularly. (B) Core and transient species exhibit different species abundance distributions for the Hinkley Point fish assemblage (Magurran and Henderson 2003). (C) Four contiguous quadrats in which species (different shapes) may be core (shaded) or transient (open). (D) The species-area relationships for (C) depending on whether transient species are excluded or not, using the lower right panel to represent the smallest area. Because every species is core in at least one quadrat, species richness at the largest scale is the same for the two relationships. (E) Temporal turnover (the Jaccard index of dissimilarity) is much lower when transient species are excluded from the calculation, since they are the species most driving turnover.

Transient species are expected to interact with their biotic and abiotic environments differently than core species since by definition they do not maintain viable local populations and are not necessarily well adapted to the local environments in which they are found (Magurran and Henderson 2003; Coyle et al. 2013; Umaña et al. 2017). Previous studies found that core species presence is more strongly tied to the environment and other deterministic factors, while transient presence is more strongly determined by stochastic factors (e.g., Magurran and Henderson 2003; Coyle et al. 2013; Umaña et al. 2017). Because much of the ecological theory related to species coexistence, niche partitioning, and biodiversity assumes that species directly interact and occur only in suitable environments, the presence of these transient species has the potential to skew our understanding of ecological systems. Indeed, transient species have been shown to differ from core species with respect to the shape of species abundance distributions (Figure 1B; Magurran and Henderson 2003), the relative importance of density-dependence versus environmental stochasticity (Magurran and Henderson 2003; Ulrich and Ollik 2004), the primary drivers of species richness (Coyle et al. 2013), and life history traits (Supp et al. 2015). We expect transient species may influence the slope of species-area relationships, since species that are transient may make up a disproportionate fraction of the community at smaller spatial scales, while at large scales most species are expected to maintain persistent populations over at least some subset of the domain (Figures 1C, D). Transient species are also likely to contribute disproportionately to estimates of temporal turnover since by definition they are present in only a small proportion of samples over time (Figure 1E). Thus, a wide variety of classic ecological patterns may differ depending on whether transient species are considered, including biodiversity patterns that have the potential to influence conservation and management decisions.

Given the potential impact of transient species on understanding and managing ecological systems, it is important to understand more about how common transient species are and how their prevalence varies with taxonomic group, ecosystem types, environmental context, and scale. There are reasons to expect that several of these factors may influence the prevalence of transients. First, species from taxonomic groups with strong dispersal abilities like birds commonly show up in habitats and regions in which they are not expected (Grinnell 1922; Coyle et al. 2013), whereas organisms with limited dispersal should do so much less frequently. Second, assemblages located in regions of high habitat heterogeneity are expected to receive more transient individuals from adjacent habitats via mass effects (Shmida and Wilson 1985). Coyle et al. (2013) found that mountainous regions had a greater proportion of transient bird species, consistent with these predictions. Third, at small scales (e.g., below the average home range size), most organisms will only be observed occasionally and the majority will therefore be classified as transients. At large scales (e.g., an entire continent), nearly all species will maintain viable populations and be consistently observed, and almost none will be classified as transient. Understanding variation in the prevalence of transient species will improve our understanding of the factors structuring communities and help identify study systems where our understanding of ecological systems is most prone to being influenced by their presence. Making comparisons across ecosystems and taxonomic groups will require understanding the scale-dependence of transient species' prevalence because the scale at which assemblages are sampled can vary by several orders of magnitude.

Here, we undertake the first systematic evaluation of the prevalence and predictors of transient species in ecological communities. We use data from over 17,000 community time series from terrestrial, aquatic, and marine ecosystems across seven major taxonomic groups to: 1) evaluate the prevalence of transient species and how it varies with taxonomic group, ecosystem type, and habitat heterogeneity; 2) assess the scale-dependence of transient species prevalence and correct for scale to make consistent comparisons across groups; and 3) examine how the inclusion of transient species in community-level analyses impacts four commonly analyzed ecological patterns including the shape of species-abundance distributions, drivers of species richness, species-area relationships, and temporal turnover.

## Methods

### Data

We conducted an extensive search for datasets of community composition over time both online and published in the literature. We identified datasets using a combination of existing compilations (Dornelas et al. 2014; Yenni et al. 2016), searching online data catalogs such as the Ecological Data Wiki (ecologicaldata.org, White 2016), exploring datasets available from Long-Ecological Research sites, exploring datasets in the data journal Ecological Archives, and conducting literature searches. We initially identified 330 datasets spanning seven broad taxonomic groups. We filtered these datasets to those meeting the following criteria: 1) each assemblage was sampled on at least six occasions (typically years, but occasionally for smaller organisms like plankton samples were monthly or bimonthly), 2) at least ten species were observed over the course of the study, and 3) the study had a spatially well-defined location with a fixed environmental context (e.g. communities based solely on the geographic coordinates of individual organisms, as in many marine pelagic transect studies, were not included). Of the 330 datasets examined, 86 satisfied our criteria and yielded 17,921 unique assemblages spanning terrestrial, marine and freshwater ecosystems. A complete list of datasets and sources is provided in T. The majority of datasets and community time series come from terrestrial bird and plant assemblages, with fewer datasets from marine and freshwater systems (Figure 2A-D). The duration of the studies ranged from six to 57 years and assemblage richness ranged from 10 to 276 species, with most assemblages having between 20 and 61 species (Figure 2E, F). All species names were checked for typos, and any taxa not identified to species (e.g. “Unidentified grass”) were removed unless the taxon clearly did not overlap with any other taxa in the dataset (e.g. “*Sigmodon* sp.” was retained only if no other *Sigmodon* species were present in the region). For datasets with uneven sampling in either space or time (e.g. variable numbers of surveys per year, or variable numbers of spatial units per survey), we standardized the level of spatial or temporal subsampling for that site in each year of the time series (see details in Appendix, Figure A1).

**Figure 2.**
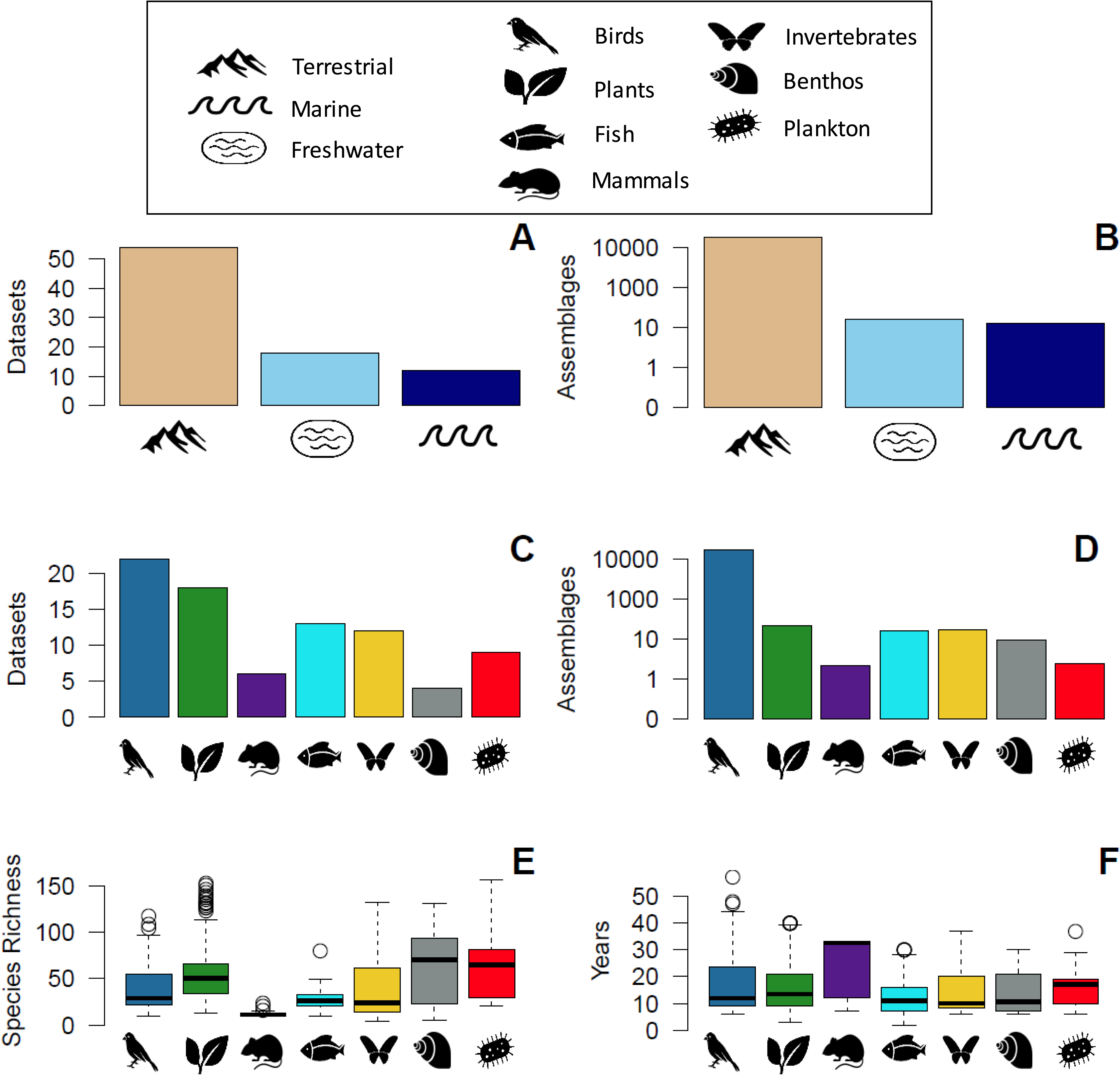
Description of the compiled time-series datasets. The number of (A) datasets and (B) number of assemblages (log scaled) by ecosystem type (terrestrial, freshwater, marine) and by taxonomic group (C, D). (E) Boxplots of the number of species per assemblage by taxonomic group. Several high richness outliers for plant and plankton assemblages were excluded to improve visualizing the bulk of the data (*). (F) Boxplots of time series length by taxonomic group.

### Analysis

Following Coyle et al. (2013), we operationally defined a species as transient at a site if it was observed in 33% or fewer of the temporal sampling intervals, and assessed the prevalence of transients as the proportion of species in the assemblage below this threshold (Figure 1A). We also evaluated more restrictive definitions using maximum temporal occupancy thresholds of 10% and 25% to evaluate the impact of this decision. Results were qualitatively similar for the three different thresholds (Figure A2-A6).

Although many authors have used the bimodality of temporal occupancy distributions (e.g., Figure 1A) to identify transient species in this way (Magurran and Henderson 2003; Dolan et al. 2009; Coyle et al. 2013), some species will be incorrectly classified due to imperfect detectability. Species with low detectability due to low density or traits or behaviors that make them difficult to detect may be persistent at a site but only detected in a small proportion of samples (MacKenzie et al. 2006). As such, estimates of the proportion of transient species based on observed temporal occupancy are likely higher than the true numbers. A full exploration of the detailed influence of imperfect detection is beyond the scope of this paper, but we are developing simulation-based approaches to understand precisely how it influences estimates of the proportion of transients as well as the identification of individual species (Hurlbert unpublished data).

While imperfect detection is clearly a concern for analyses of this type there is also evidence that using observed occupancy provides a reasonable first approximation of transient status. Magurran and Henderson (2003) showed that using occupancy to identify species as transient is consistent with using habitat preferences. In an examination of nearly 500 bird communities, Coyle et al. (2013) showed that transient species richness was correlated with regional habitat heterogeneity as would be expected of true transients while it was not positively correlated with vegetation which would be expected to impede species detections. In addition, similar studies using habitat preference-based transient designations (Belmaker 2009) have yielded similar conclusions to those using occupancy based approaches (Coyle et al. 2013). Finally, the results in this paper are similar for species that are comprehensively surveyed and those that are less thoroughly sampled (see *Results* and *Considerations*). So, while there is no doubt that misclassifications will occur, for large data compilations like this one that lack both detailed habitat preference data for species and the necessary sampling methods to estimate detection probabilities, occupancy based approaches appear to provide a reasonable approximate classification. We address these issues further in the *Considerations* section of the Discussion.

We evaluated the effect of spatial scale on the perceived prevalence of transient species using the subset of datasets that included sampling at hierarchically nested spatial scales. We used a linear mixed model to quantify how the proportion of transient species in an assemblage varied with the spatial scale over which the assemblage was characterized. The model included taxonomic group as a fixed effect and dataset as a random effect, with both variables having the potential to influence both the slope and intercept of the relationship. Area was log-transformed for analysis. Because scale will be perceived differently for organisms of different size—e.g. a 1 ha quadrat is effectively much larger for ants than for birds—it may not allow for direct comparisons of “scale” among taxonomic groups. As such, we also built a similar mixed model using the median community size for all assemblages (i.e., the total number of individuals sampled in an assemblage, median = 102) as an alternative, potentially more generalizable, measure of scale.

To explore the influence of habitat heterogeneity on the prevalence of transients we used a linear mixed model to predict the proportion of transients as a function of elevational heterogeneity (the variance in elevation within a 5 km radius of the site), with spatial scale (using community size as a proxy) as a covariate and taxonomic group as a random effect. P-values were estimated from the *t-*statistics using a normal approximation. All terrestrial datasets with geographic coordinates were used to fit the model. We used a 30 arc-second digital elevation model DEM of North America (GTOPO30), acquired from the USGS Earth Resources Observation and Science Center (EROS), to calculate the variance of elevation. We calculated a pseudo R^2^ for each mixed model based on the fit between predicted and observed values.

Finally, we quantified the influence of transient species on a suite of commonly studied ecological patterns including species-abundance distributions, species-area relationships, temporal turnover, and correlates of species richness. We did this by comparing the form of these patterns when using data on the entire community to the same pattern generated after excluding species that were identified as transients (i.e. those species with temporal occupancy ≤ 33%). We fit two distributions for species-abundance, the logseries and the Poisson lognormal to the combined abundance data across years for each time-series. Magurran and Henderson (2003) proposed that transient species should be better fit by the logseries and core species by the lognormal, meaning that excluding transient species should result in improved fits by the lognormal. We compared the fits of the two distributions based on AIC_c_ model weights. Analysis of species-area relationships was restricted to datasets with hierarchical spatial sampling. Power function relationships were fit to each assemblage using linear regression on log-transformed data (Xiao et al. 2011) to predict the number of species observed from the area sampled. The fitted exponents of the relationships were compared. Mean temporal turnover was calculated as the mean of the Jaccard dissimilarity index (Krebs 1999; Figure 1E) between all adjacent time samples in each community time series. Analyses of the drivers of species richness were restricted to data from the Breeding Bird Survey of North America since it was the only dataset that employed consistent sampling across large spatial scales with a large number of replicates. For this last set of analyses we used two environmental correlates that are known to be important for determining richness in this dataset, the Normalized Difference Vegetation Index (NDVI), a remotely sensed estimate of productivity, and elevation (White and Hurlbert 2010). We calculated correlation coefficients between each environmental variable and species richness (including or excluding transient species), as well as correlation coefficients for transient species richness alone to further illuminate differences.

The complete set of R scripts for data cleaning and processing are available on Github (http://www.github.com/hurlbertlab/core-transient) and analysis scripts for this study are archiveat Data Dryad (URL to be filled in).

## Results

Assemblages from all ecosystem types and taxonomic groups included a substantial proportion of transient species, and relatively few species with intermediate temporal occupancies (Figure 3). The proportion of an assemblage made up of transient species varied with taxonomic group, with means ranging from 32-58%. Benthos and invertebrates had more than 50% of species characterized as transient on average. Fish, plankton, and plant communities had 46-49% transient species on average. In mammal communities, 45% of species were classified as transient, while birds had the lowest proportion of species classified as transients at 30%. Terrestrial ecosystems had the lowest proportion of transient species (37%) followed by marine (48%) and freshwater (55%) systems (Figure 3B).

**Figure 3.**
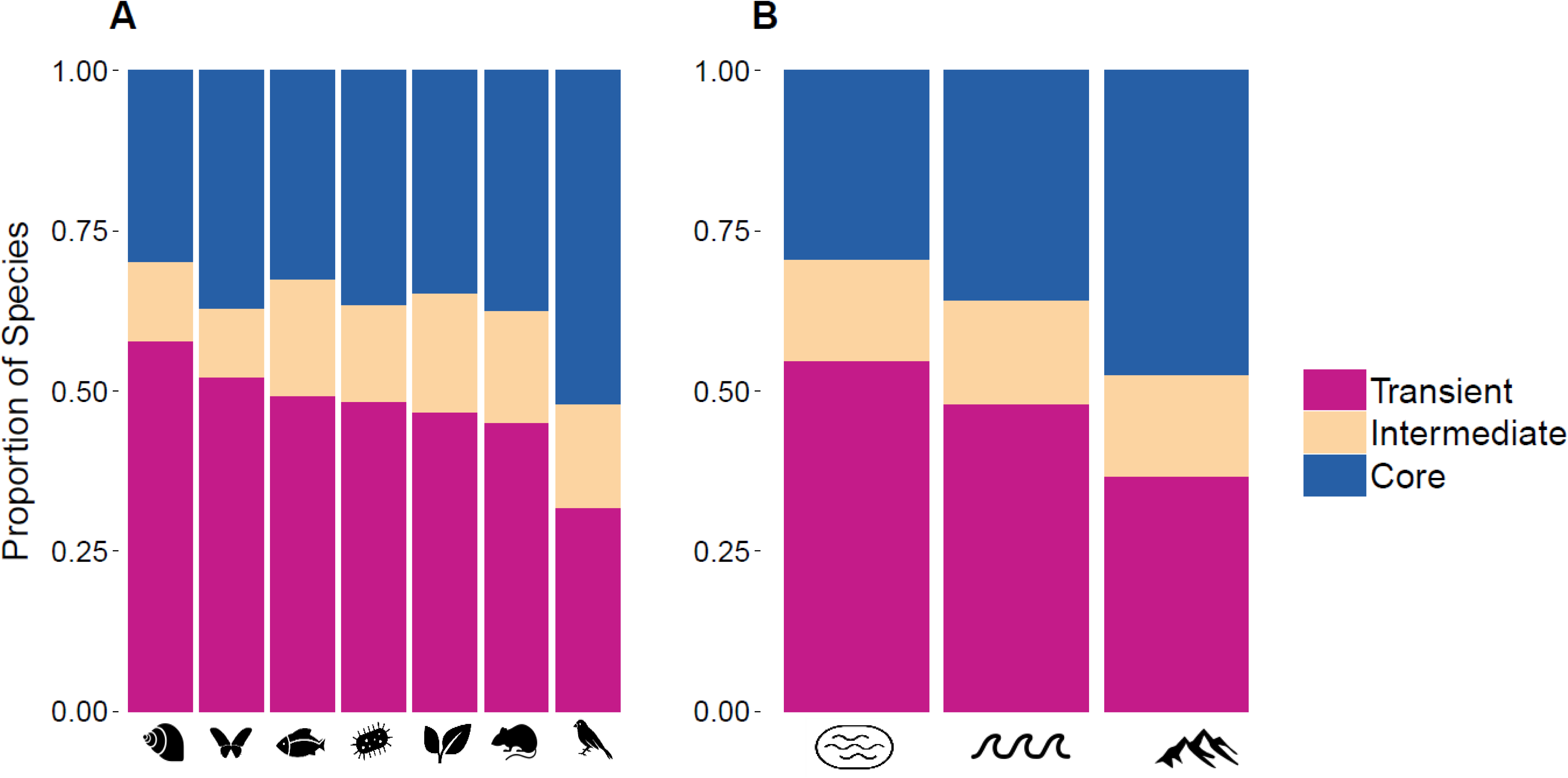
The mean proportion of species in an assemblage that are transient (≤ 33% temporal occupancy), core (>66.7%), or neither, grouped by (A) taxonomic group and (B) ecosystem. See Figure 2 for icon key.

There was a negative effect of sampling area on the proportion of transients in a community (*p* < 10^-16^), but scaling relationships varied substantially in both slope and intercept across datasets and taxonomic groups (Figure 4A; pseudo R^2^ = 0.01). When we characterized the scaling relationships using total community size based on the total number of individuals in an average sample instead of sample area the relationship was considerably stronger (Figure 4B; pseudo R^2^ = 0.32). Communities at scales in which large numbers of individuals are sampled have few transient species, while communities at scales in which small numbers of individuals are sampled have proportionally more transient species, regardless of taxonomic group. After controlling for scale (community size), birds—one of the taxonomic groups with the lowest representation of transient species based on the raw survey data—became comparable to benthic and terrestrial invertebrates, which had the highest representations of transient species based on raw data (cf. Figure 3A and 4C). Mammal and plankton communities had the lowest average proportion of transient species in scale-corrected datasets at approximately 40%. Controlling for sampling scale, the proportion of transients in an assemblage no longer varied across type of ecosystem (Figure 4D).

**Figure 4.**
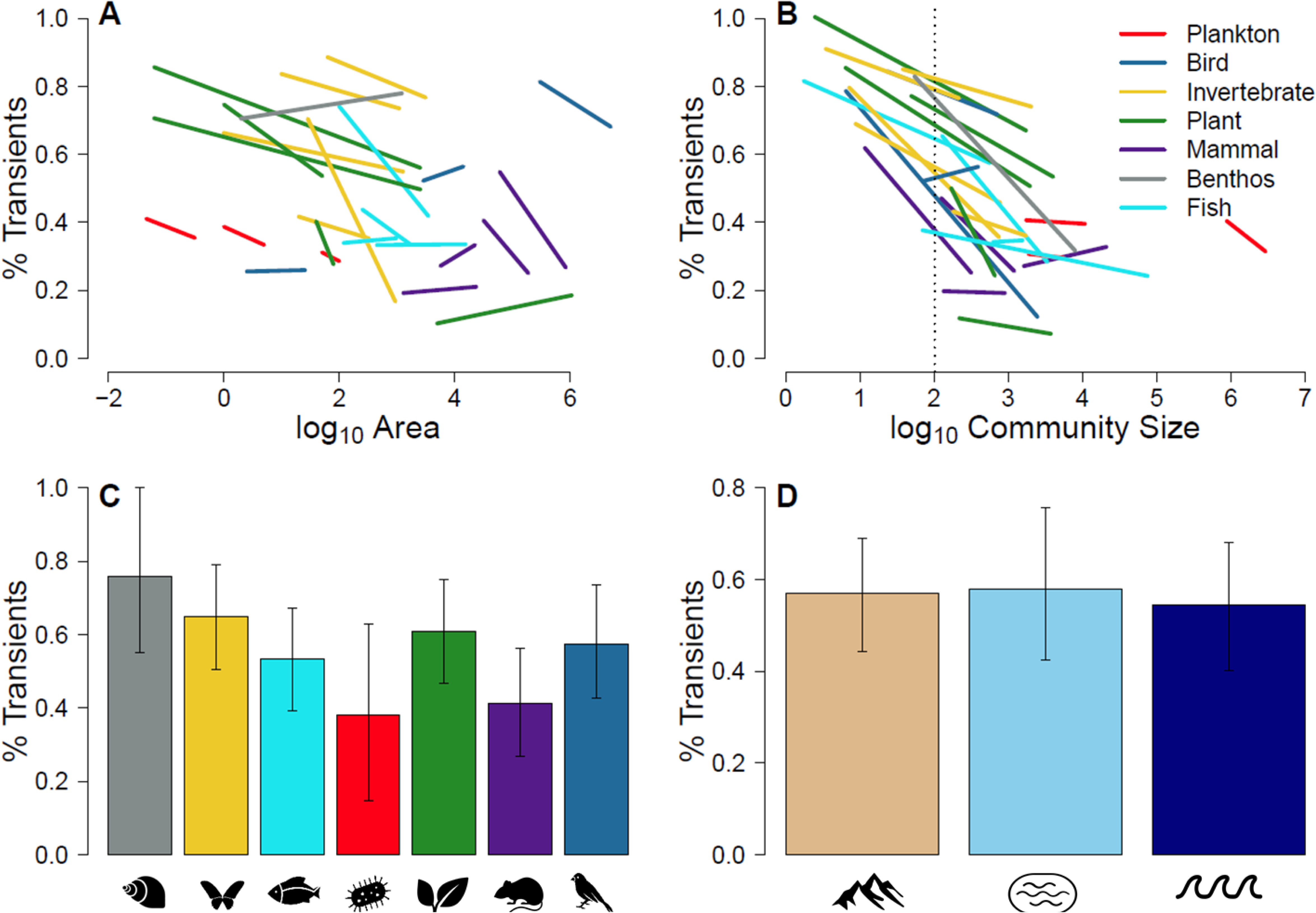
Linear models of the proportion of transient species as a function of (A) sample area and (B) sample community size (number of individuals) for each dataset with a spatially hierarchical sampling scheme. Datasets are color coded by taxonomic group. The proportion of transient species expected for a hypothetical community of 102 individuals (the median community size across datasets) for a given (C) taxonomic group or (D) ecosystem based on linear mixed effects models (see text). No spatially hierarchical datasets were available to evaluate benthic invertebrates. See Figure 2 for icon key.

Elevational heterogeneity was found to have a positive effect (*p* <0.0001) on the proportion of transient species when accounting for community size as a covariate and taxonomic group as a random effect (Table 1). There was no evidence for an interaction between elevational heterogeneity and community size (*p* = 0.98).

**Table 1.**
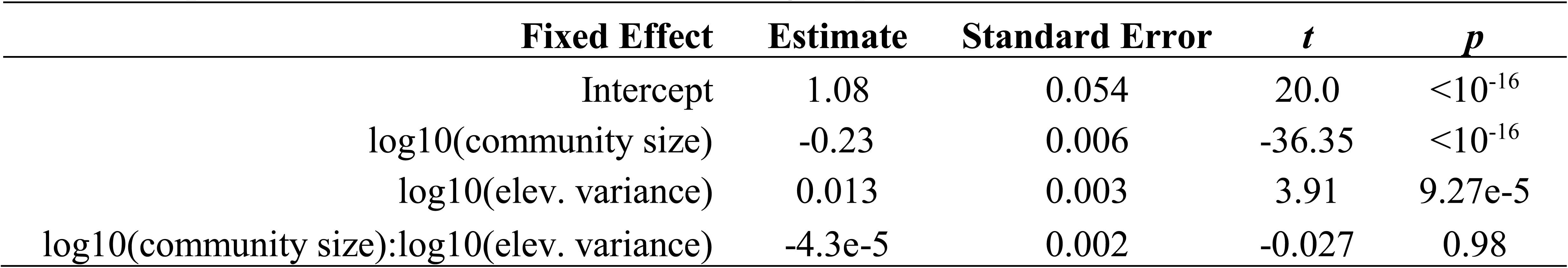
Linear mixed model results for the effect of elevational heterogeneity and community size on the proportion of transients. Taxonomic group was included as a random effect.

| Fixed Effect | Estimate | Standard Error | *t* | *p* |
| --- | --- | --- | --- | --- |
| Intercept | 1.08 | 0.054 | 20.0 | <10<sup>-16</sup> |
| log10(community size) | -0.23 | 0.006 | -36.35 | <10<sup>-16</sup> |
| log10(elev. variance) | 0.013 | 0.003 | 3.91 | 9.27e-5 |
| log10(community size):log10(elev. variance) | -4.3e-5 | 0.002 | -0.027 | 0.98 |

Finally, we examined whether the inclusion of transient species impacted four fundamental ecological patterns. Species abundance distributions for full assemblages were generally best fit by a logseries distribution, although there was some support for the lognormal, whereas assemblages excluding transient species were universally better fit by a lognormal distribution (Figure 5A). The strength of species richness drivers varied depending on whether transient species were included or not, because transient species exhibited environmental correlations of opposite sign to non-transient species (Figure 5B). As such, excluding transient species led to a stronger positive correlation between richness and the vegetation index NDVI, (0.53 versus 0.48), and a stronger negative correlation with mean elevation (−0.46 versus −0.37). Species turnover was always higher when transient species were included than when they were excluded, with an average deviation of 0.11 (Figure 5C). Finally, the exponent of the species-area relationship was typically higher when excluding transients (average deviation = 0.07; Figure 5D). All results were similar using alternative occupancy thresholds to define transient species (Figures A2-6).

**Figure 5.**
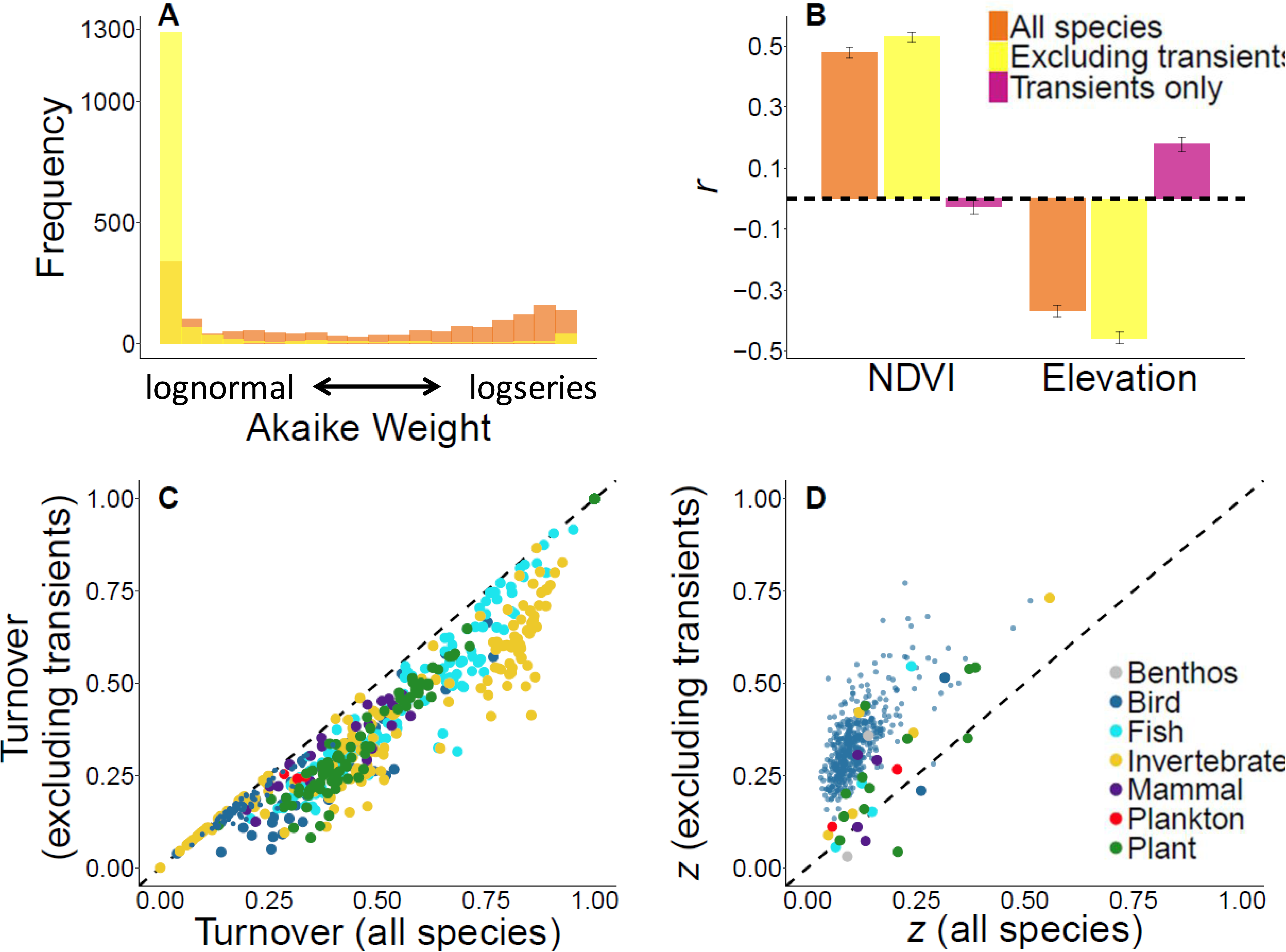
Comparison of common ecological patterns between full communities and communities excluding transient species. (A) Histogram of Akaike weights for the logseries model of the species abundance distribution for all species (orange) and excluding transients (yellow). Because only two models were compared, Akaike weights close to 0 imply strong support for the lognormal model. (B) Environmental correlates of species richness (NDVI and elevation) including transients (orange), excluding transients (yellow), and transients only (pink). (C) Comparison of temporal turnover estimates when including or excluding transient species. Temporal turnover was quantified using the Jaccard dissimilarity index. Points are color coded by taxa and small blue circles represent the North American Breeding Bird Survey. (D) Comparison of species-area relationship exponents when including or excluding transient species. Points are the same as in (C).

### Discussion

We quantified the prevalence and impact of transient species in ecological communities using data on over 17,000 community time series spanning multiple ecosystems, taxonomic groups, and spatial scales. Transient species were a common element of communities in all taxa and ecosystems examined, demonstrating that these species are a general feature of ecological systems. Transient species interact with their abiotic and biotic environment in distinct ways (Magurran and Henderson 2003; Ulrich and Ollik 2004; Coyle et al. 2013; Umaña et al. 2017), which highlights the need to better understand the contexts in which transient species are expected to be prevalent and the potential impact transient species may have on ecological inferences.

The largest source of variation in the proportion of transient species observed in a community is related to spatial scale. For communities sampled at multiple spatial scales, the proportion of transient species decreased with increasing scale, as species were more likely to be observed and actually persist over larger sampling areas. As a result, comparisons of the prevalence of transient species between studies should account for scale. However, area per se may not be directly comparable between communities that differ substantially in body size or otherwise use space differently. An alternative measure of scale, community size, effectively controls for differences in area usage between taxonomic groups by integrating the influence of each species’ distinct life history traits and home range sizes. Correcting for scale in this way, we found that the proportion of transient species did not vary with ecosystem type, whereas ignoring scale would have led to the conclusion that transient species were much more common in freshwater than terrestrial ecosystems. Similarly, correcting for scale led to a more even distribution of the proportion of transient species across taxonomic groups, and some groups that would otherwise have been inferred to differ substantially in the prevalence of transients were actually found to be comparable.

Differences in the prevalence of transient species were evident among taxonomic groups even when controlling for spatial scale. Invertebrate, plant, and bird communities had the highest proportion of transient species while plankton and mammal communities had the lowest. These taxonomic groups differ in many respects precluding a rigorous analysis, but we speculate that traits such as dispersal ability and habitat specialization may increase the likelihood of species being temporarily observed in areas where they are not well adapted and hence being recorded as transients. For example, birds have strong dispersal ability relative to the other taxonomic groups and there are numerous records of individuals spotted far outside their geographic range and in unexpected habitats (Grinnell 1922). Similarly, plants with passive seed dispersal may be transported great distances and may consequently be more likely to be observed in unsuitable habitat (Willson 1993). Small mammals have more limited dispersal, which may explain why mammal communities (dominated in our dataset by small mammal communities) have a lower proportion of transient species on average. The plankton datasets examined in this study came primarily from lakes, and low rates of dispersal between lakes could explain the low proportion of transient species for this group.

In addition to dispersal, groups composed of more generalist species might be expected to have a lower proportion of transient species because most species can maintain viable populations in most locations. The low prevalence of transient species in plankton communities may also be explained by this phenomenon, as Hutchinson (1961) noted “paradoxically” that most plankton species are generalists that compete for the same limited resources. Specialist species, on the other hand, will only maintain populations in select locations with suitable conditions, allowing mass effects (Shmida and Wilson 1985) or accidental dispersal to result in transient occurrences in other areas.

In addition to trait differences among taxa, variability in the prevalence of transient species was related to environmental heterogeneity. Transient species were more prevalent in communities with higher elevational heterogeneity, which extends the findings of Coyle et al. (2013) for birds to a broader range of taxa. Homogeneous landscapes tend to have homogeneous communities (Stegen et al. 2013; Stein et al. 2014) and a site within such a landscape is unlikely to receive immigrants from poorly adapted species compared to a site in a heterogeneous landscape with a more diverse species pool from more diverse habitats. Indeed, environmental heterogeneity and species richness are frequently positively related (Stein et al. 2014), and our results indicate this may be due in part to an increase in transient species rather than an increase in habitat specialists (Gaston et al. 2007; Stein et al. 2014).

### Impacts of Transient Species on Ecological Inference

The presence of transient species in ecological communities influenced all of the ecological patterns we examined, from measures of local community structure, to spatial and temporal turnover, to richness gradients at continental scales. This highlights the importance of considering transients when trying to manage and understand ecological communities. The species abundance distribution (SAD) characterizes the relative abundance of common and rare species in communities and different distributions have been associated with different processes structuring the community (McGill et al. 2007; Connolly et al. 2014). Building on the results of Magurran and Henderson (2003), we show that including transient species in an analysis results in more logseries-like SADs while excluding them results in more lognormal distributions. This result is consistent with the idea that different processes influence the community assembly of transient versus core species (Henderson and Magurran 2014; Supp et al. 2015). Based on theoretical grounds, many SAD models may be more appropriately applied to all species observed, or only to the set of species that strongly interact and maintain viable populations. For example, neutral theory applies to all species, as it explicitly allows for rare immigration or speciation events (Hubbell 2001), whereas resource allocation based niche apportionment models (MacArthur 1957; Tokeshi 1990) are likely more appropriately applied only to non-transient species. While the SAD may not be sufficient on its own to infer community structuring processes (Cohen 1968; Volkov et al. 2005; Baldridge et al. 2016; but see Connolly et al. 2014), it is one of several ecological patterns that may collectively shed light on such mechanisms (McGill et al. 2007; Blonder et al. 2014). As such, consideration of transient species has the potential to influence our understanding of local community structure.

In addition to influencing measures of local community structure, the inclusion of transient species also affected measures of how ecological systems turnover and change with scale. Estimates of temporal turnover were always higher when transients were included in assemblages. This occurs because transient species are only present over a small fraction of a time series, resulting in higher turnover in species composition within a community over time (see also Magurran and Henderson 2010). Conversely, the inclusion of transient species led to lower estimates of spatial turnover as reflected in the slope of species-area relationships. This is because a greater proportion of the species list at small spatial scales is identified as transient compared to at a larger scale. As such, including transient species increases richness more at small scales than large, resulting in a shallower species-area relationship and lower spatial turnover (Figure 1D). Turnover and associated scaling relationships have implications for assessment of community responses to global change (Brown et al. 1997; Suding et al. 2008), understanding processes structuring spatiotemporal variation in communities (Adler et al. 2005; McGlinn and Palmer 2009), and up and downscaling biodiversity estimates for conservation (Shen and He 2008; Azaele et al. 2015; Kitzes and Harte 2015), further indicating that consideration of transients is important for understanding local to regional scale ecological systems.

Finally, inclusion of transient species also influenced the strength of continental scale correlates of species richness. Excluding transient species increased the explanatory power of both NDVI and elevational heterogeneity. Transient species correlations were opposite of those observed for core species, consistent with our general findings on the relationships between environment and heterogeneity (Coyle et al. 2013). Because the proportion of transient species varies along environmental gradients, analyses at large scales will potentially weight core and transient species differently in different locations and the perceived importance of environmental associations with ecological patterns may often change when excluding transient species. In this example, the inclusion of transients weakens the perceived support for a species-energy relationship (Wright 1983; Hurlbert 2004) compared to when only non-transients were considered. Given the impact on a wide range of ecological patterns, the decision to include or exclude transient species in a community analysis is an important one that should be made by explicitly considering the nature of the conceptual framework or theory being investigated. In some cases, it will be necessary to remove these species from analyses or risk making improper inferences.

### Considerations

Conceptually, transient species are those that do not maintain persistent populations over time and therefore only appear infrequently during surveys. The bimodality of temporal occupancy distributions (e.g., Figure 1A) has led many authors to suggest that temporal occupancy can be used to distinguish these transient species from more core members of a community. However, it can be difficult to tease apart whether species of low occupancy are truly transient or simply have low density or detectability (Henderson and Magurran 2014). We followed Coyle et al. (2013) in using a maximum occupancy threshold of 33% as our operational definition of transient species, but all of the results we report here were similar using stricter thresholds of 10% or 25% (Figures A2-A6). If the focus were on a single community, then the accuracy of identifying transient species might be improved by a combination of assessing the shape and natural break points of each community's particular occupancy distribution, and incorporating information on species habitat preference as done by Belmaker (2009) for coral reef fish. Alternatively, when the sampling design allows for the estimation of detection probabilities it should be possible to correct for these issues using occupancy modeling (MacKenzie et al. 2006). Independent validation of transient status (e.g., by evidence of breeding, or knowledge of habitat affinities) or occupancy modeling based approaches are always desirable when possible, and analyses along environmental gradients should carefully consider how detectability might vary along such gradients (Coyle et al. 2013). However, for many groups detailed information on habitat preferences or estimates of true population persistence is not readily available, and a definition based on a universal occupancy threshold is currently the most feasible option for analyzing hundreds or thousands of assemblages for cross-taxon comparisons like those presented here.

As described in the *Methods*, there is evidence that occupancy based thresholds provide reasonable identifications of transient species (Magurran and Henderson 2003; Belmaker 2009; Coyle et al. 2013). There is additional evidence from our results that using this raw occupancy based approach provides a reasonable approximate classification. First, the misclassification rate should presumably be lower when defining transient species using stricter occupancy thresholds, and so the consistency of our results across multiple thresholds lends some confidence to this approach. Second, for certain communities the taxonomic group and mode of data collection provide nearly complete censuses of all individuals within a static sample (e.g. plant stems within a quadrat or fish in a seine net). In these communities, imperfect detection should have little influence on estimates of occupancy (at the scale of sampling). The similarity of results in this study across groups that tend to be thoroughly surveyed (e.g., plants and fish) and those that are less intensively sampled (e.g., birds and butterflies) suggests that our results are not driven heavily my misclassifying imperfectly detected species. A detailed understanding of when and to what extent imperfect detection probabilities influence the assessment of the prevalence and impact of transient species will require simulation based approaches (Hurlbert unpublished data).

### Conclusions

Our results show that transient species are prevalent in ecological communities across all taxa, scales, and ecosystems examined. Despite the ubiquity of these species, most studies in community ecology have implicitly ignored this concept by characterizing communities using surveys that provide a snapshot of community composition in time. Because transient species interact with their biotic and abiotic environment differently—in most cases, more weakly—than non-transient species, their inclusion in community analyses impacts a wide range of ecological patterns including estimates of community structure, turnover, and biodiversity. Ecologists should explicitly consider whether to include or exclude transient species in analyses by determining whether the theories, conceptual frameworks, and conservation interests of their research are best aligned with entire communities including transient species or with only core species that maintain sustained populations at a site. A failure to do so may result in inappropriate tests of models, incorrect inferences regarding processes, and imperfect conservation efforts. When data are unavailable for distinguishing species in a community as transient or not, researchers should be aware of how this uncertainty may bias their results. While some methodological challenges remain, future studies will benefit from considering when and how the inclusion of transient species impacts our understanding of how communities respond to environmental gradients, habitat fragmentation, climatic shifts, and other disturbances.

## Acknowledgements

This research was supported the National Science Foundation through grant DEB-1354563 to A.H. Hurlbert and E.P. White and by the Gordon and Betty Moore Foundation's Data-Driven Discovery Initiative through Grant GBMF4563 to E.P. White. Taxa and ecosystem symbols are from the nounproject.com and are credited as follows: bird - parkjisun, plant - Delwar Hossain, mammal - Francisca Arévalo, fish - Iconic, butterfly - Jacqueline Fernandes, snail - B Barrett, plankton - Boris Belov, mountain - Alice Noir, waves - Alex Muravev, lake - Anton Gajdosik. The following datasets requested specific acknowledgements. Toolik LTER: This material is based upon work supported by the National Science Foundation under Grants #DEB-1026843, 981022, 9211775, 8702328; #OPP-9911278, 9911681, 9732281, 9615411, 9615563, 9615942, 9615949, 9400722, 9415411, 9318529; #BSR 9019055, 8806635, 8507493. Any opinions, findings, conclusions, or recommendations expressed in the material are those of the author(s) and do not necessarily reflect the views of the National Science Foundation. Kellogg Biological Station: Support for this research was also provided by the NSF Long-term Ecological Research Program (DEB 1637653) at the Kellogg Biological Station and by Michigan State University AgBioResearch. Cedar Creek: This work was supported by grants from the US National Science Foundation Long-Term Ecological Research Program (LTER) including DEB-0620652 and DEB-1234162. Further support was provided by the Cedar Creek Ecosystem Science Reserve and the University of Minnesota. Konza Prairie LTER: Data for Plant species composition on selected watersheds at Konza Prairie, Konza LTER small mammals, Konza LTER grasshopper monitoring, Weekly record of bird species observed on Konza Prairie, and Fish population on selected watersheds at Konza Prairie was supported by the NSF Long Term Ecological Research Program at Konza Prairie Biological Station. Hubbard Brook: Data on the Bird Abundances at the Hubbard Brook Experimental Forest (1969-present) and on three replicate plots (1986-2000) in the White Mountain National Forest were provided by Richard Holmes on 6/16/2016. These data were gathered as part of the Hubbard Brook Ecosystem Study (HBES). The HBES is a collaborative effort at the Hubbard Brook Experimental Forest, which is operated and maintained by the USDA Forest Service, Northern Research Station, Newtown Square, PA. Significant funding for collection of these data was provided by DEB 0423259 (Hubbard Brook Long Term Ecological Research). Sevilleta LTER: Sevilleta LTER mammals data set was provided by the Sevilleta Long Term Ecological Research (LTER) Program. Significant funding for collection of these data was provided by the National Science Foundation Long Term Ecological Research program (NSF Grant numbers BSR 88-11906, DEB 9411976, DEB 0080529 and DEB 0217774). Maryland Biological Stream Survey: Data included in this document were provided by the Maryland Department of Natural Resources Monitoring and Non-tidal Assessment Division. Lake Kasumigaura: The Lake Kasumigaura database, Table 10 Phytoplankton density, Lake Kasumigaura database, Table 12-1 Density of Rotifer, Cladocera and Copepoda, Lake Kasumigaura database, Table 14-1 Benthos data, and Lake Kasumigaura database, Table 15-2 Fish density data are those of the Lake Kasumigaura Long-term Environmental Monitoring Program of the National Institute for Environmental Studies, Japan. Data collection for the icthyoplankton time series northeast of Taiwan was supported by the Council of Agriculture and the Ministry of Science and Technology, Taiwan (to CHH). We are grateful to the thousands of scientists and volunteers who have helped to collect and share all of the data analyzed herein.

## Appendix

### Sampling standardization of community time series datasets

For the most accurate estimates of temporal occupancy, the sampling intensity used to characterize the assemblage in question should be identical every year. However, for some datasets, sampling intensity was not uniform in space or in time. In some cases, the number of spatial units (e.g. plant quadrats) censused varied by sampling date, in other cases, the number of sampling dates per year with which an assemblage could be characterized varied between years, and occasionally both spatial and temporal subsampling levels varied.

We used a sample-based rarefaction approach (Gotelli 2008) to standardize the effort with which an assemblage is characterized over its time series. Choosing the number of spatial or temporal subsamples to use for rarefaction is a non-trivial problem, however. On the one hand, the lowest common level of subsampling across years or sites might be chosen which will enable the inclusion of all years or sites available in the dataset. In this case, data from the best sampled sites or years will be thrown out during rarefaction in order to compare those sites or years with the less well sampled ones. On the other hand, one might choose a high level of subsampling which will provide a more thorough characterization of those well sampled assemblages. In this case, data will be lost as many sites or years will not meet this high threshold and so will not be included in the analysis.

For each dataset, we attempted to maximize both the number of assemblages that would be available for analysis as well as the thoroughness with which an assemblage was characterized by choosing the lowest level of subsampling that was met by at least 50% of possible site-years (Figure A1).

---

**Figure A1.**
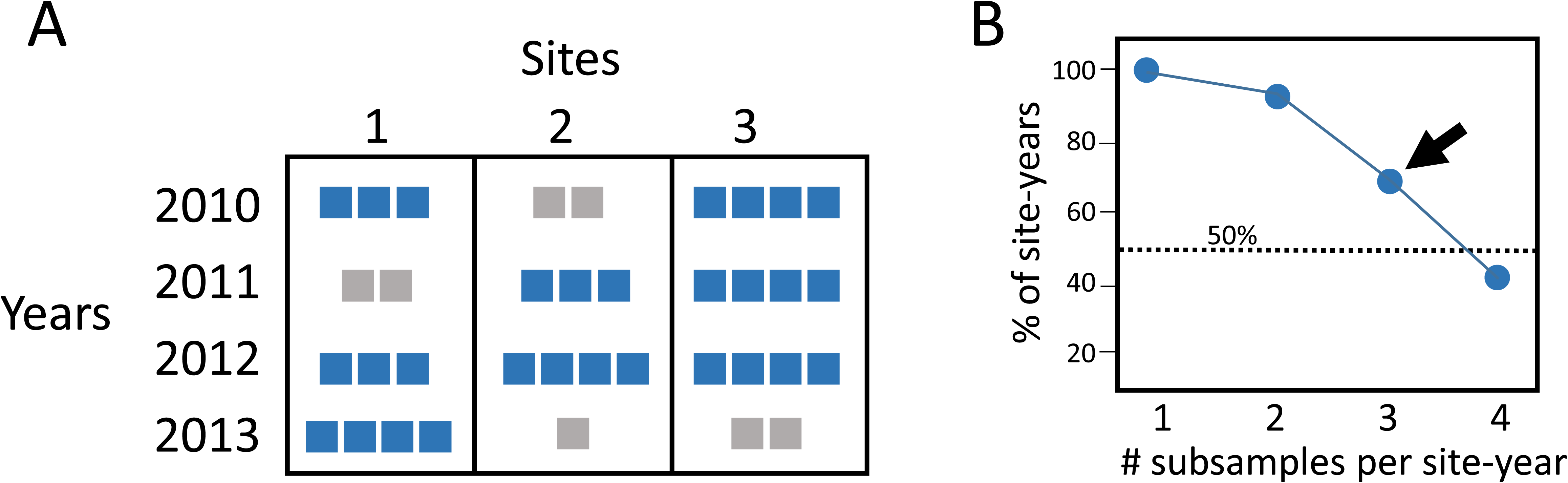
(A) A hypothetical dataset with 3 sites that have been sampled with variable intensity over time (squares). Site-years exceeding the threshold of 3 subsamples are highlighted in blue. Only data from these site-years would be used in an analysis, and where the total number of subsamples exceeds the threshold, only 3 would be chosen at random to characterize an assemblage. (B) The subsampling threshold is the smallest value for which the % of available site-years exceeds 50% (dotted line).

**Table A1.**
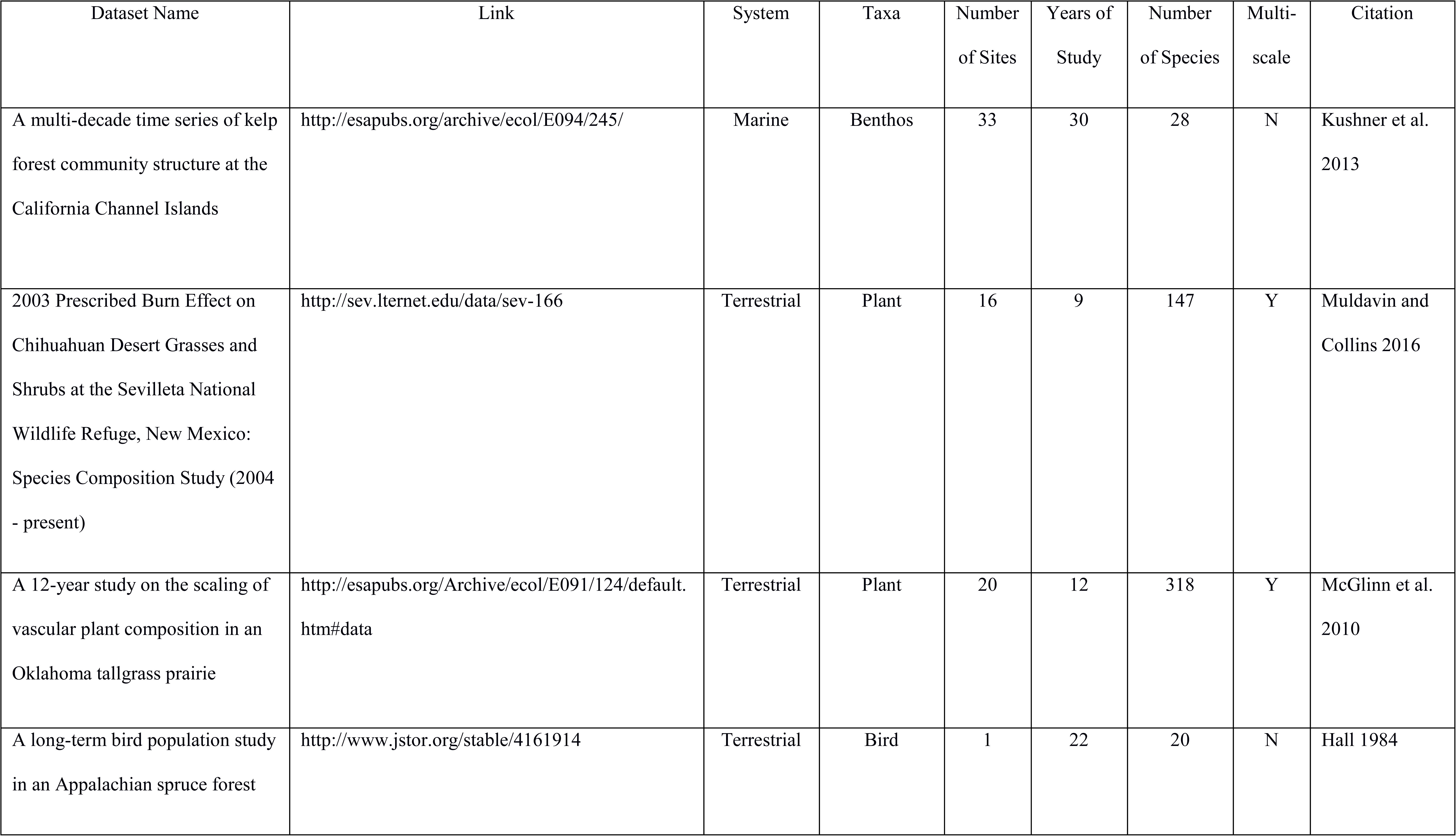
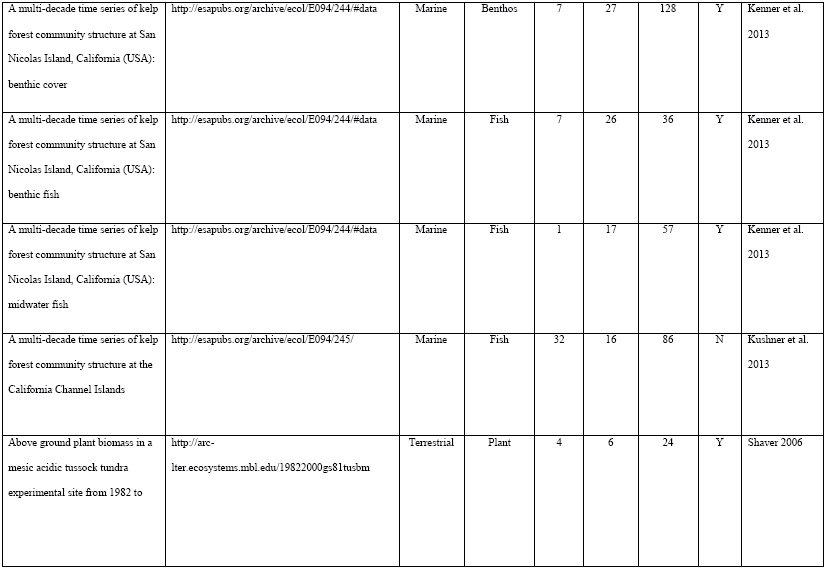
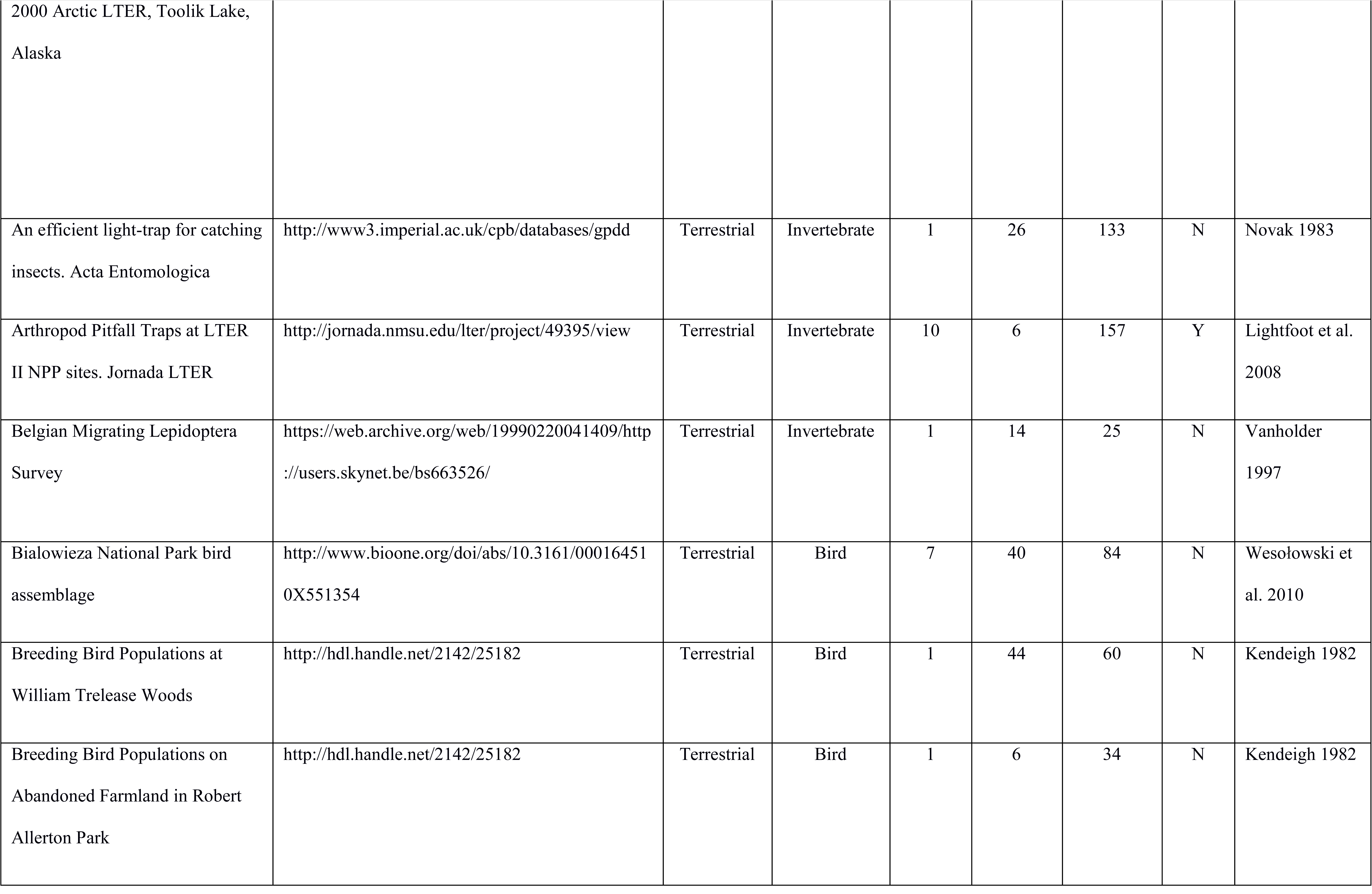
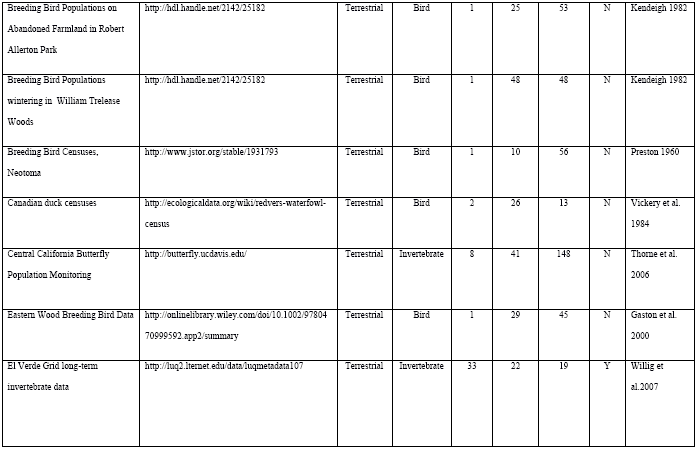
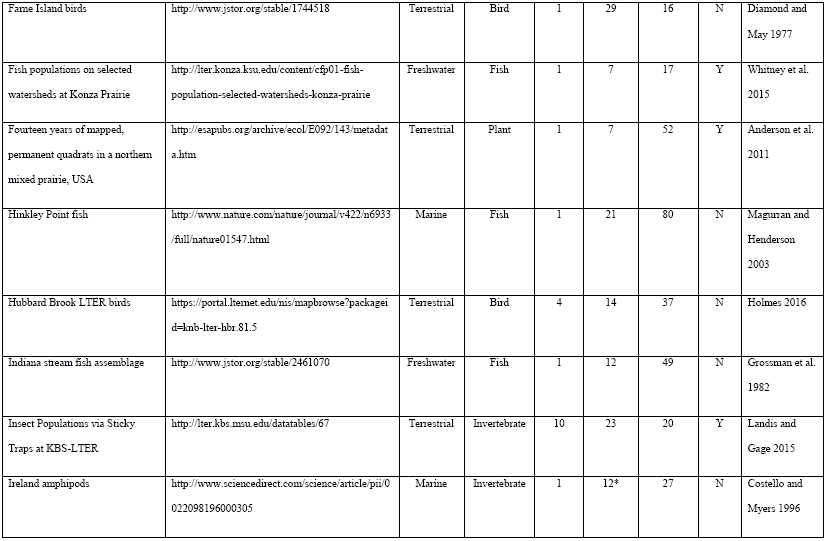
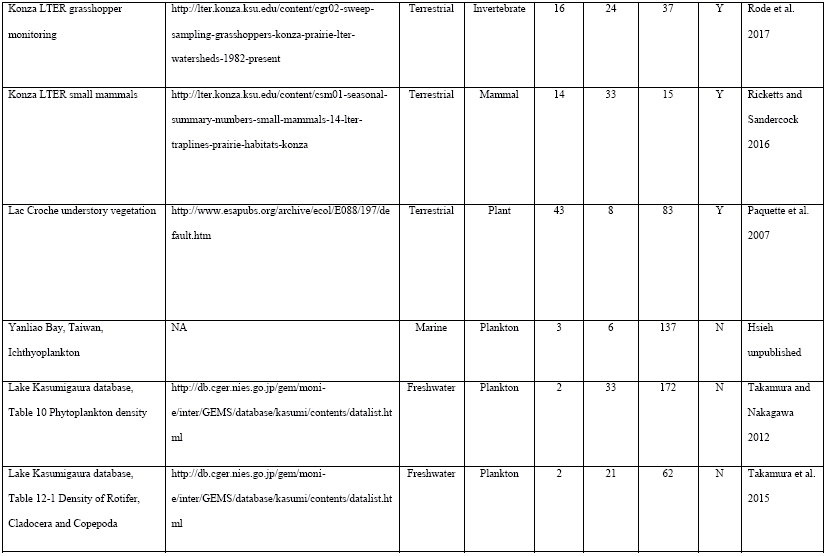
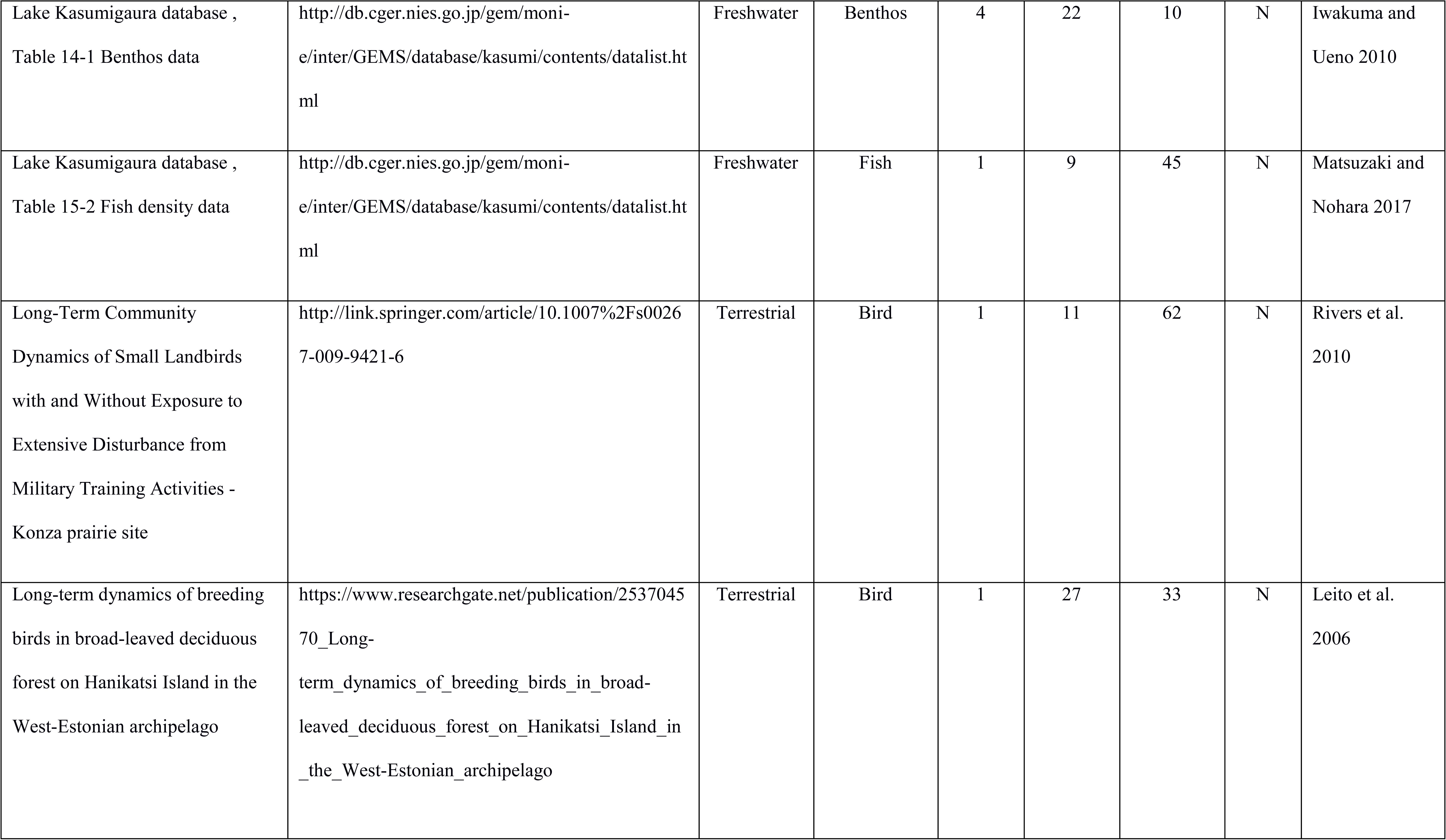
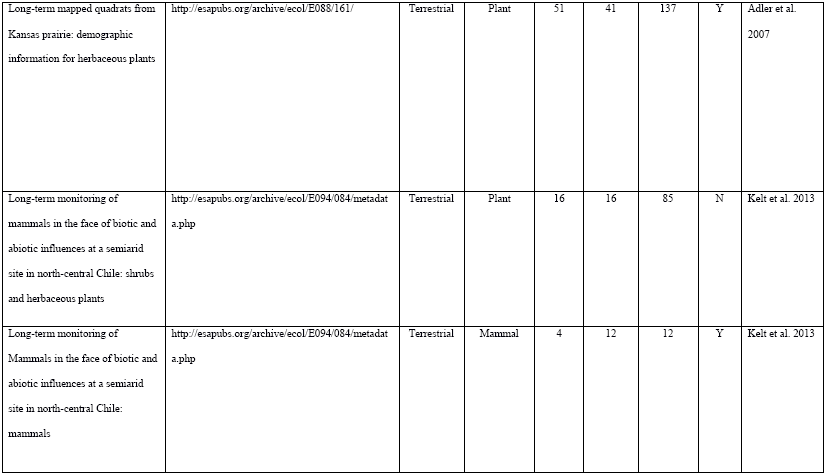
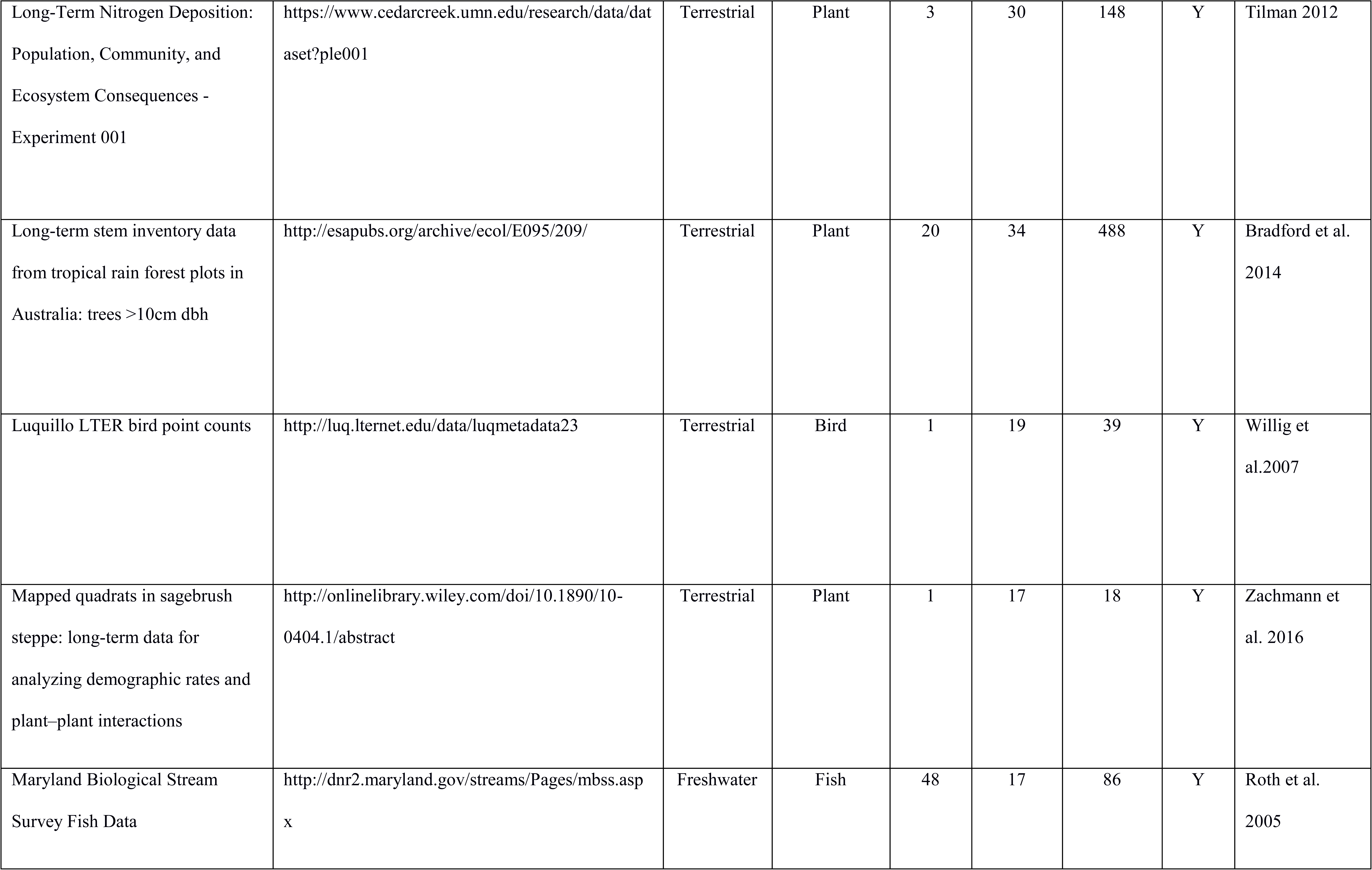
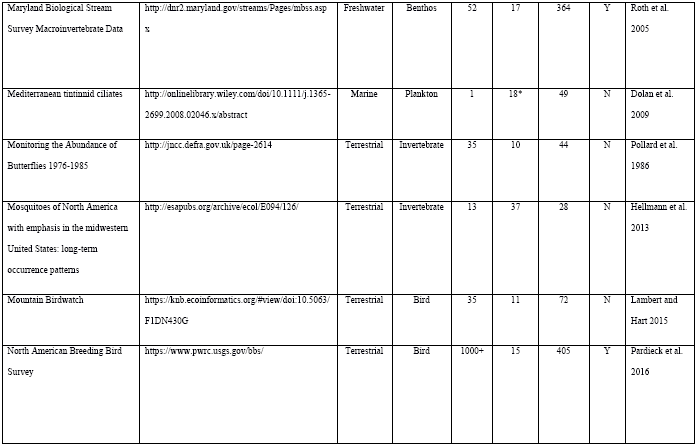
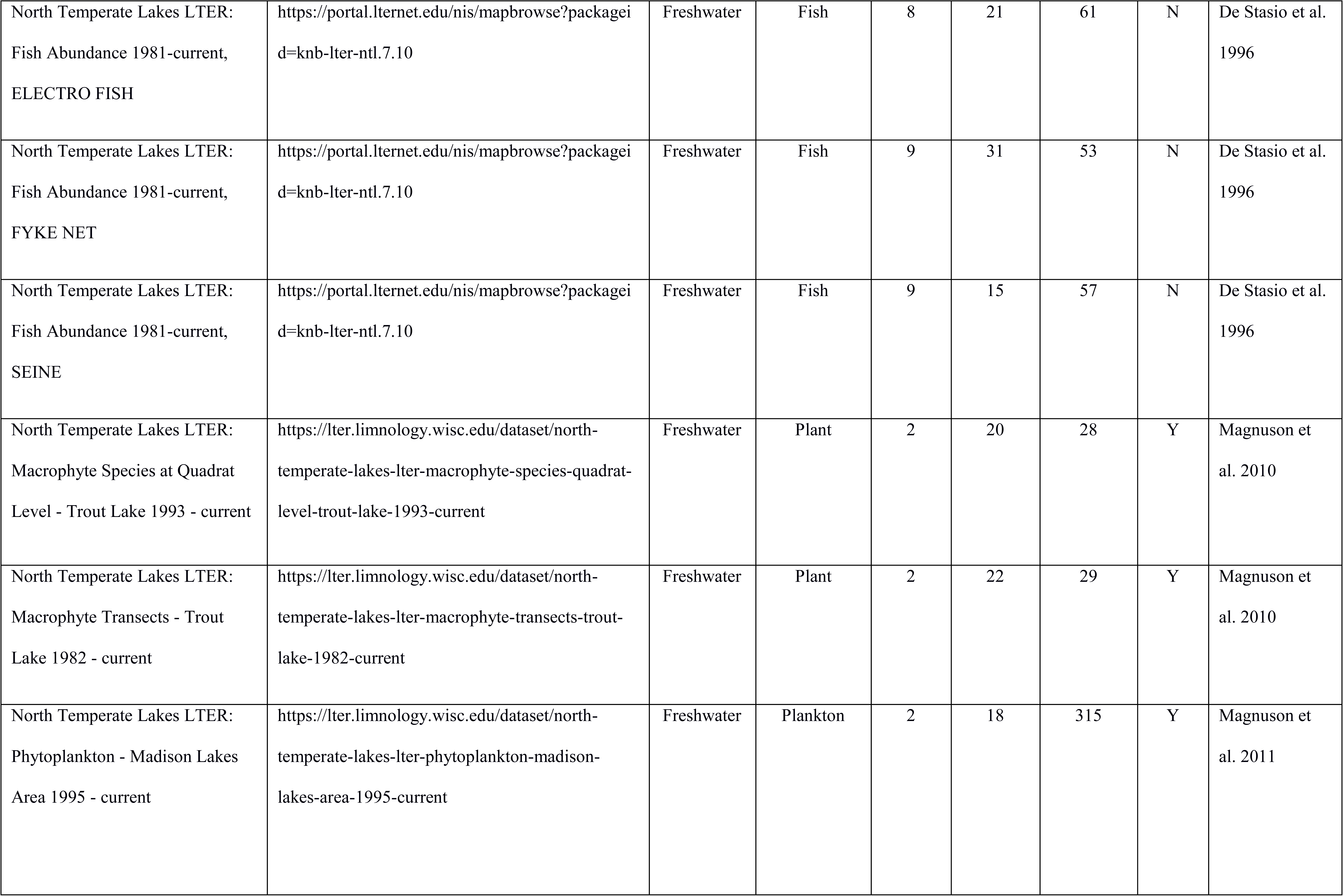
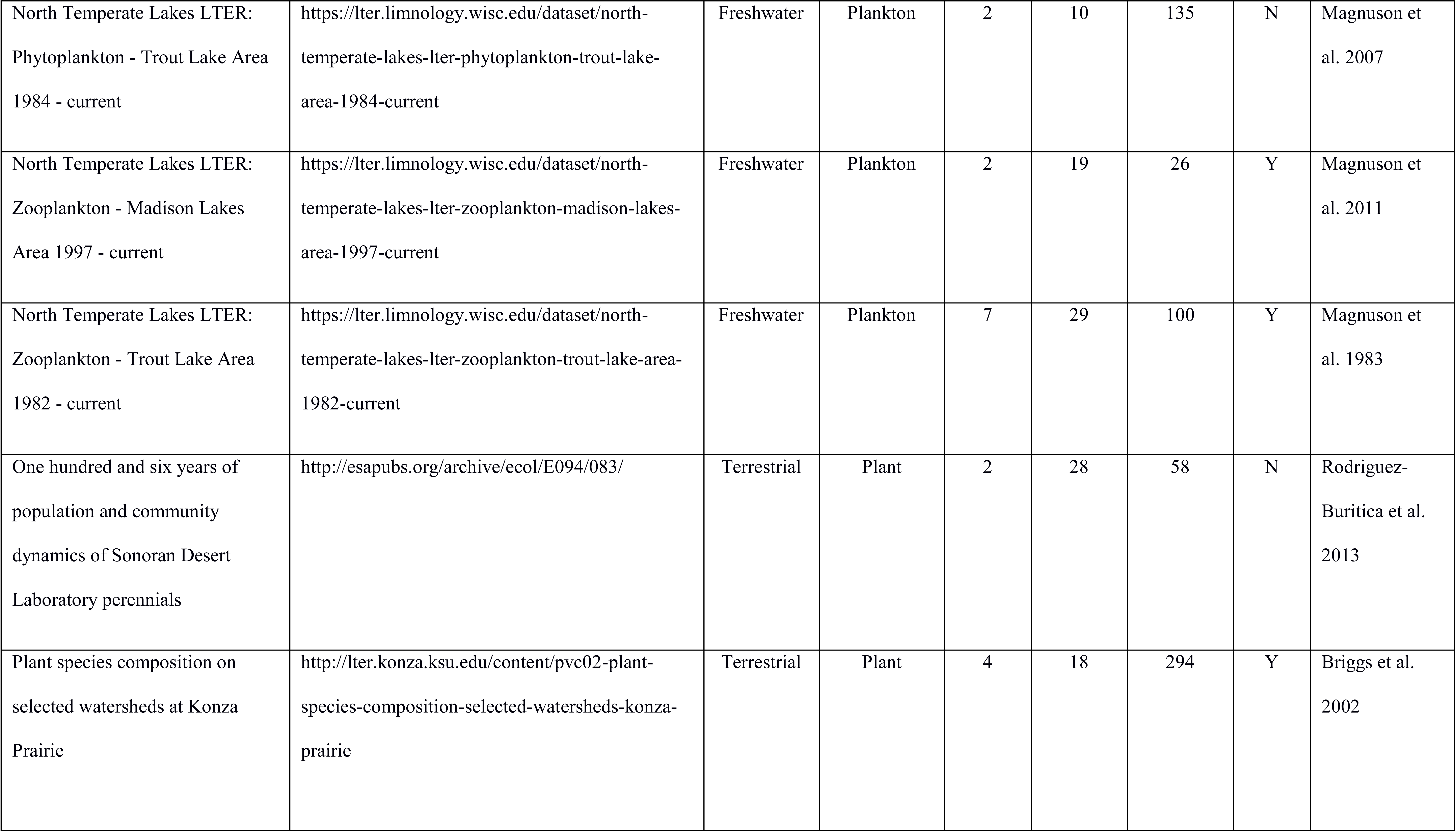
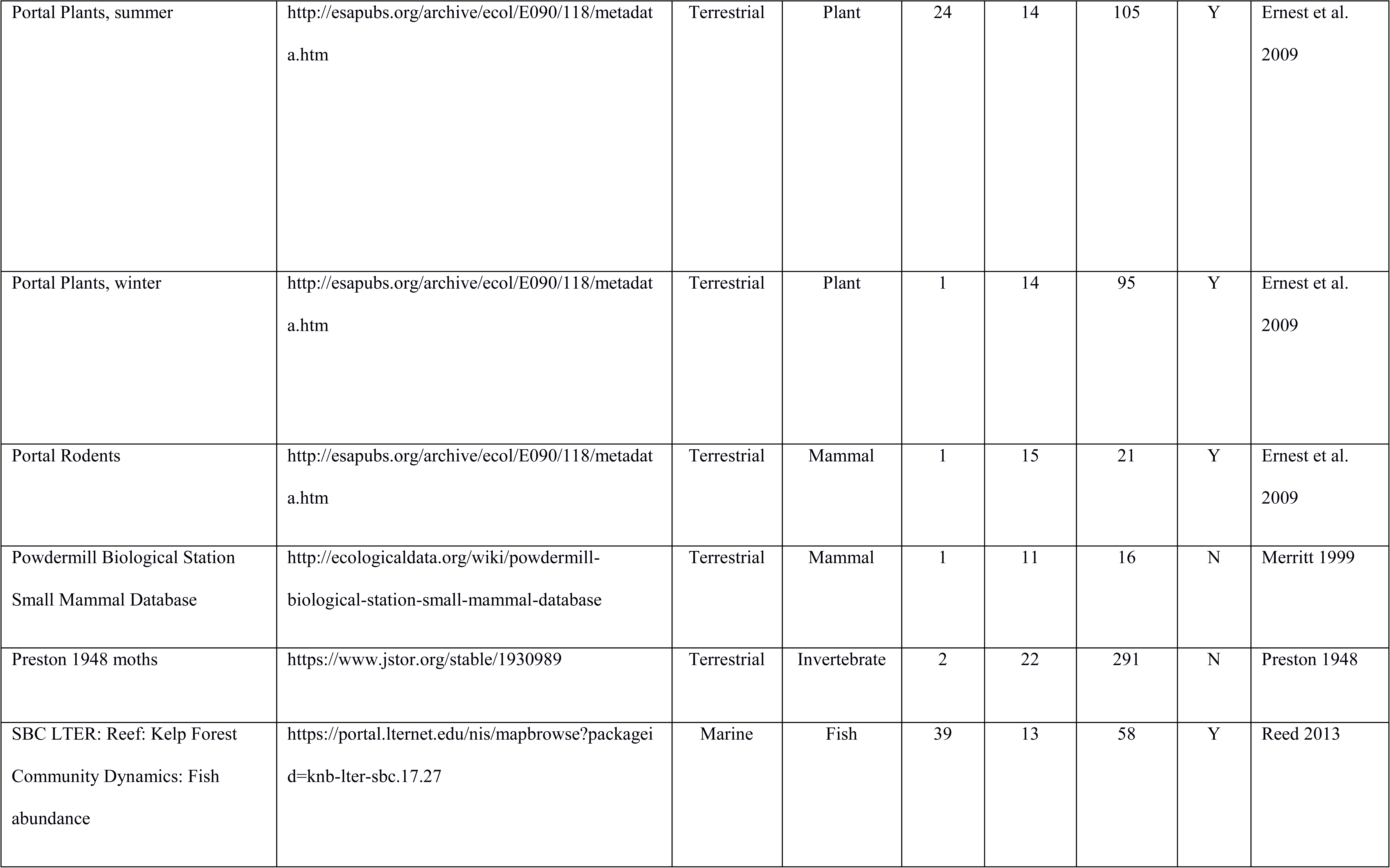
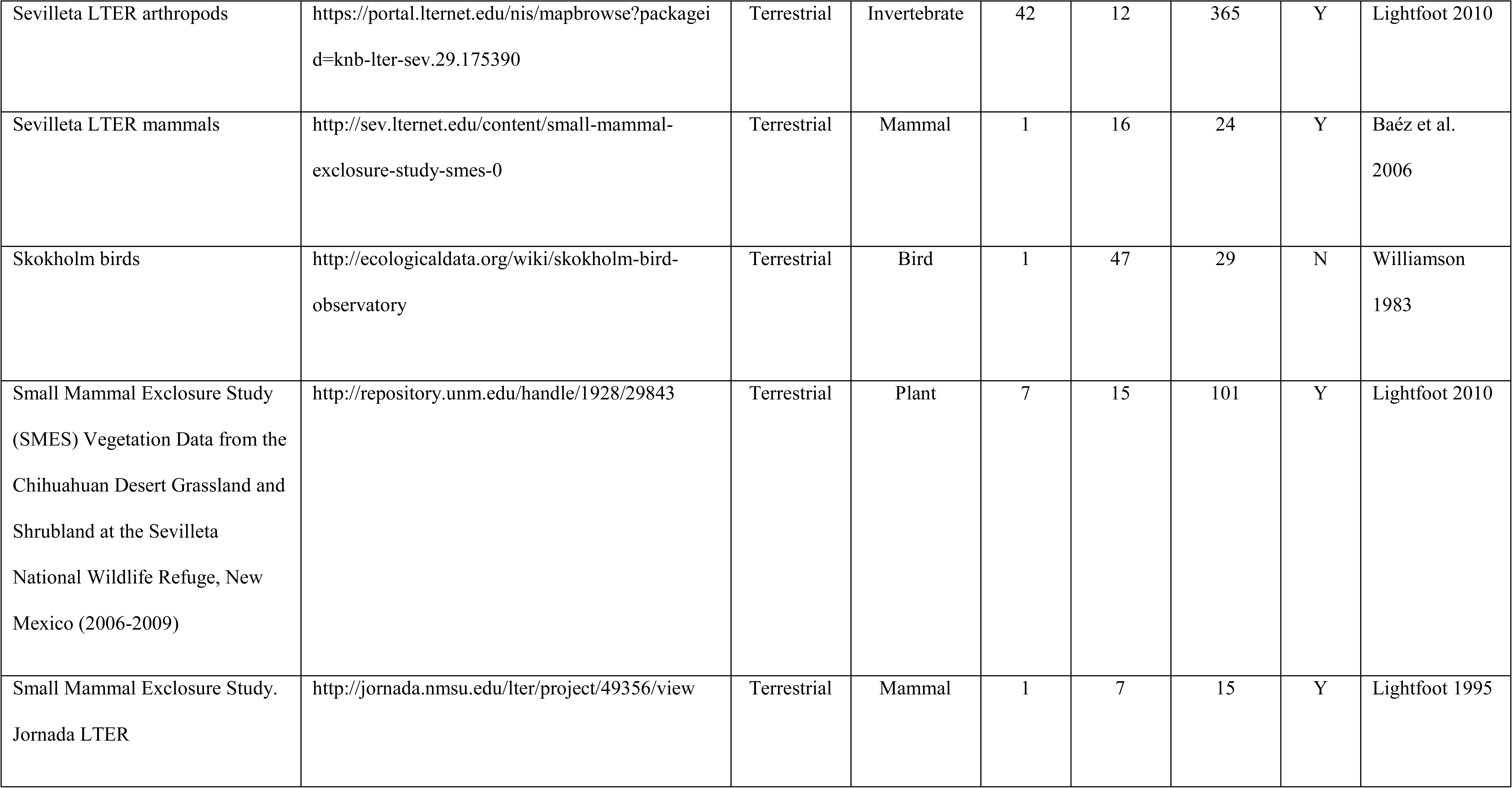
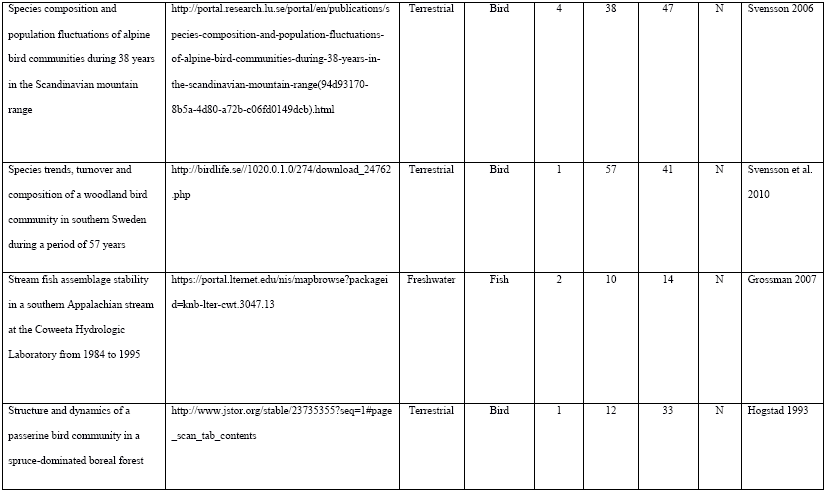
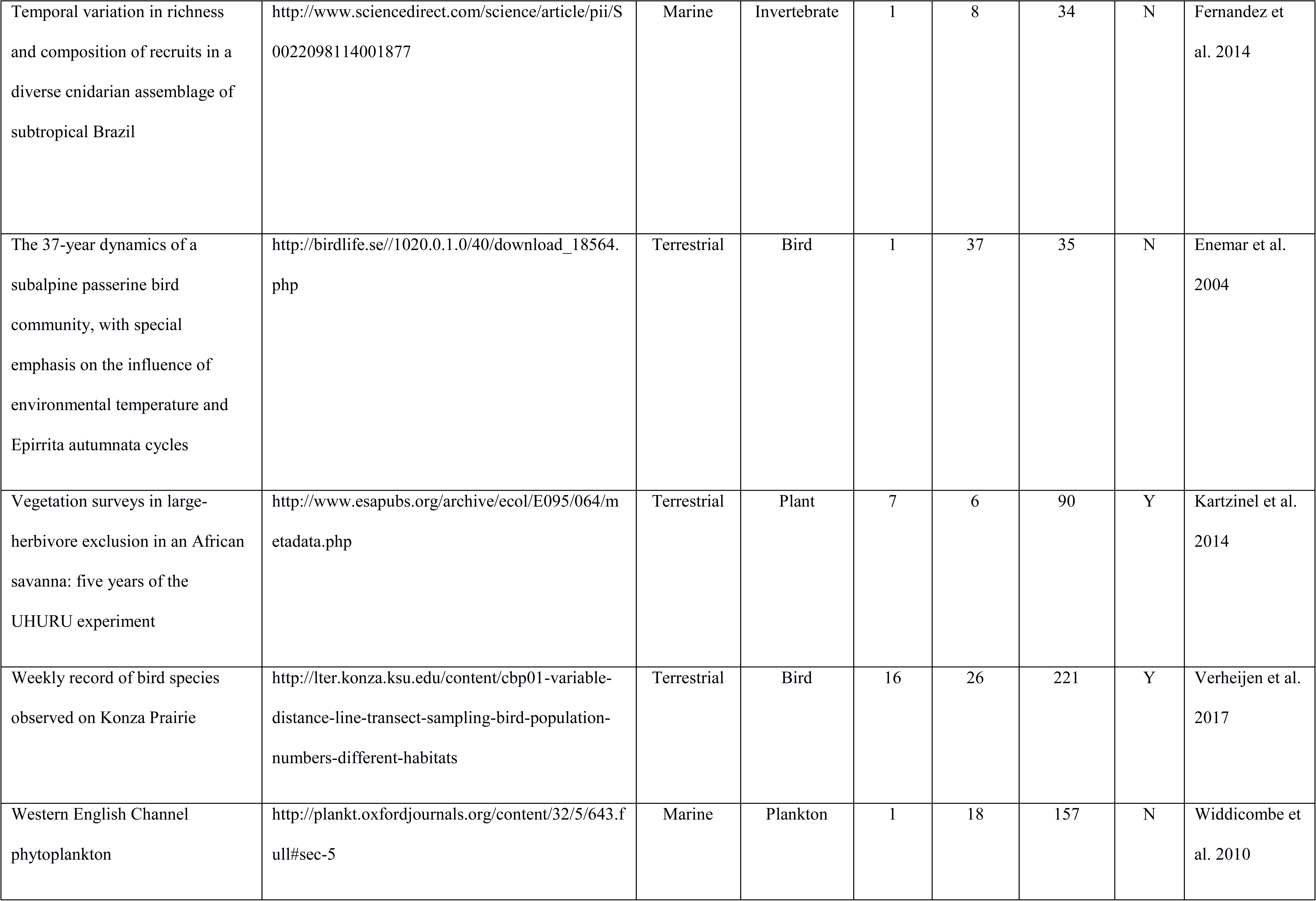
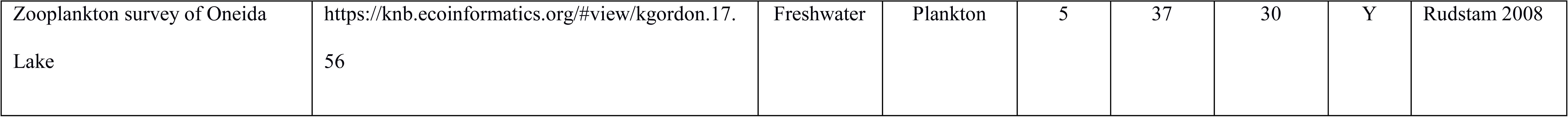
Table of datasets, sources, and citations used for analyses.

**Figure A2.**
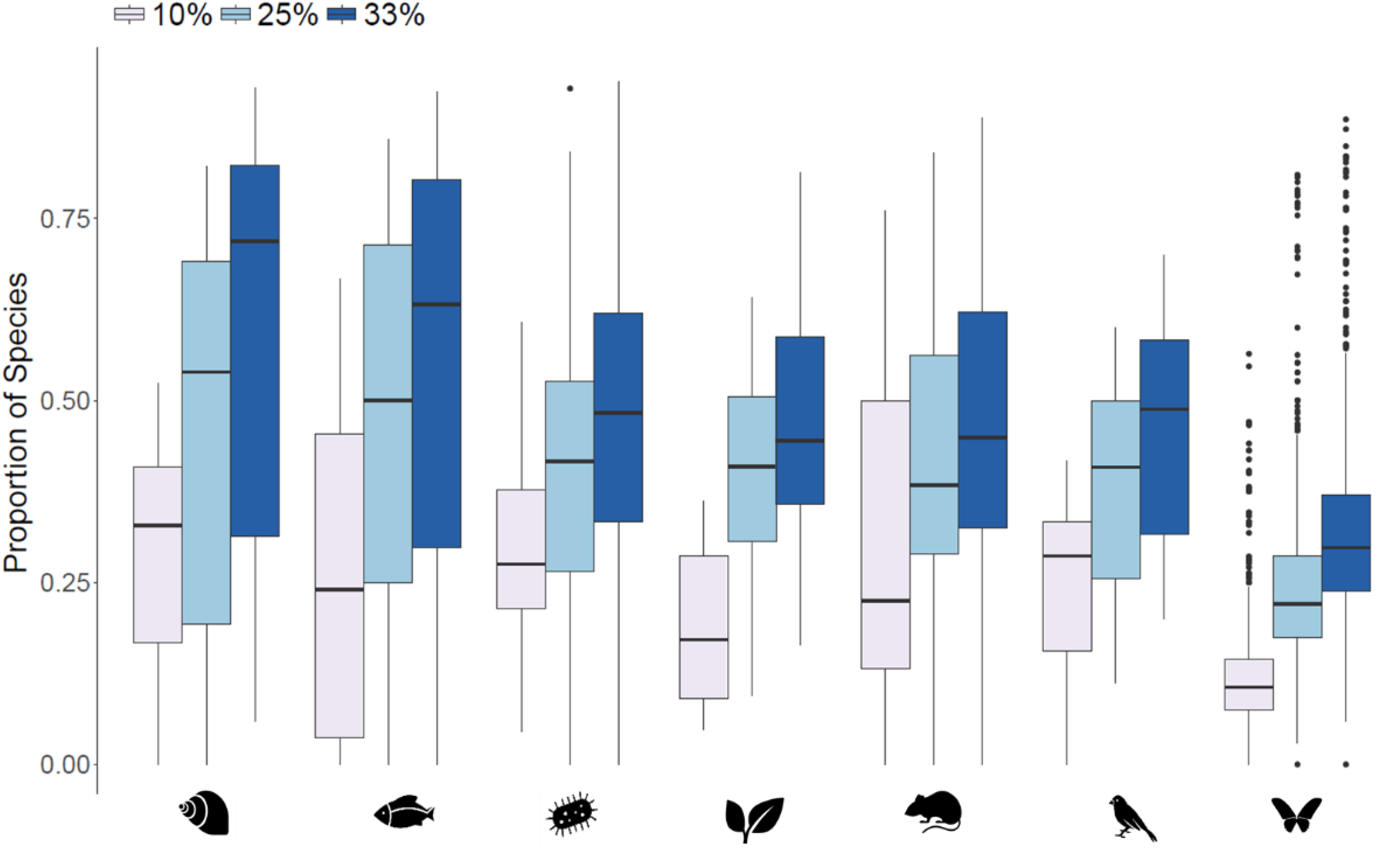
The impact of different thresholds on the proportion of transient species in assemblages from different taxonomic groups. Taxon symbols as in Figure 2.

**Figure A3.**
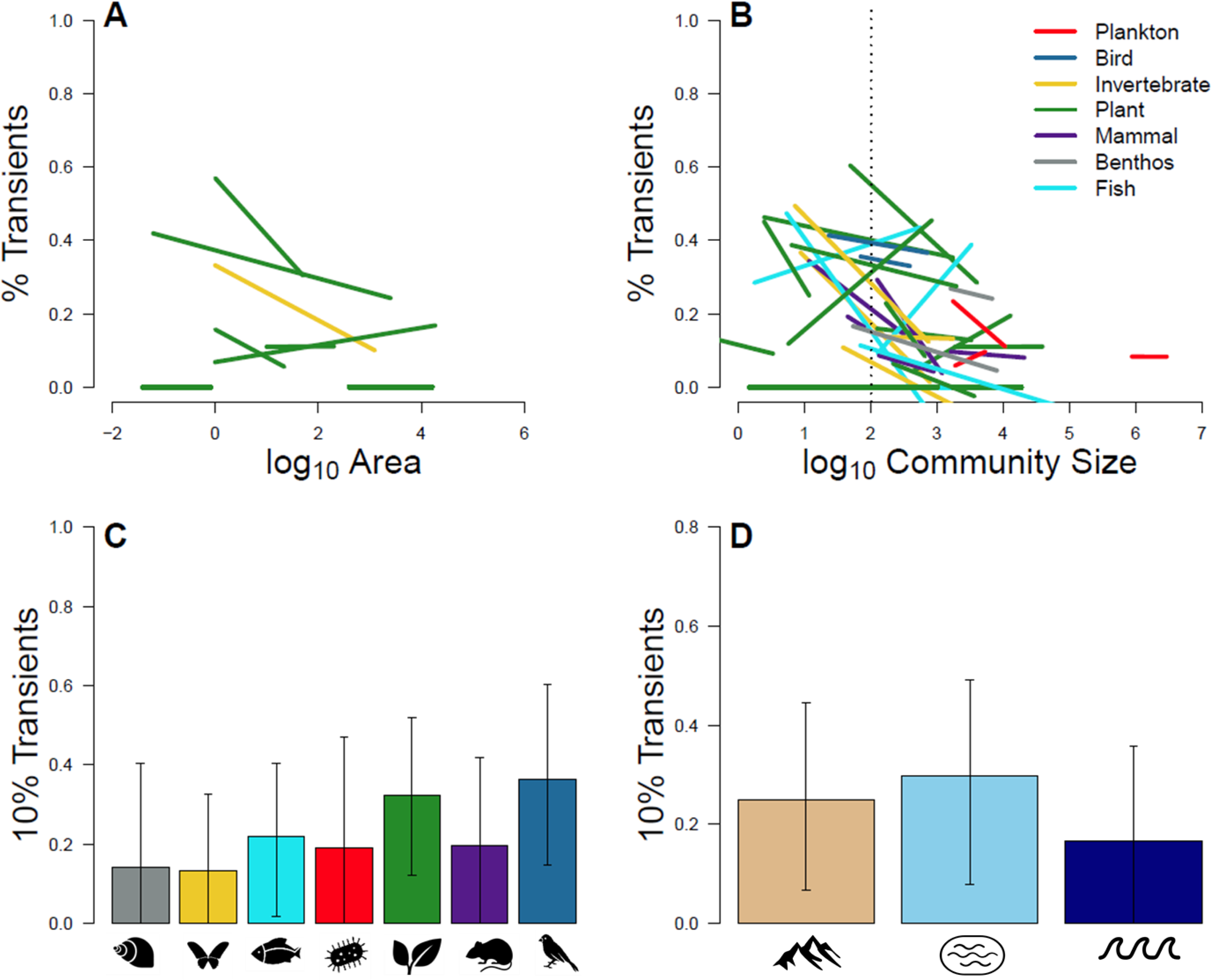
The impact of scale on the proportion of transient species as displayed in Figure 4 in the main text, but where transient species are defined as those with temporal occupancy ≤ 10%.

**Figure A4.**
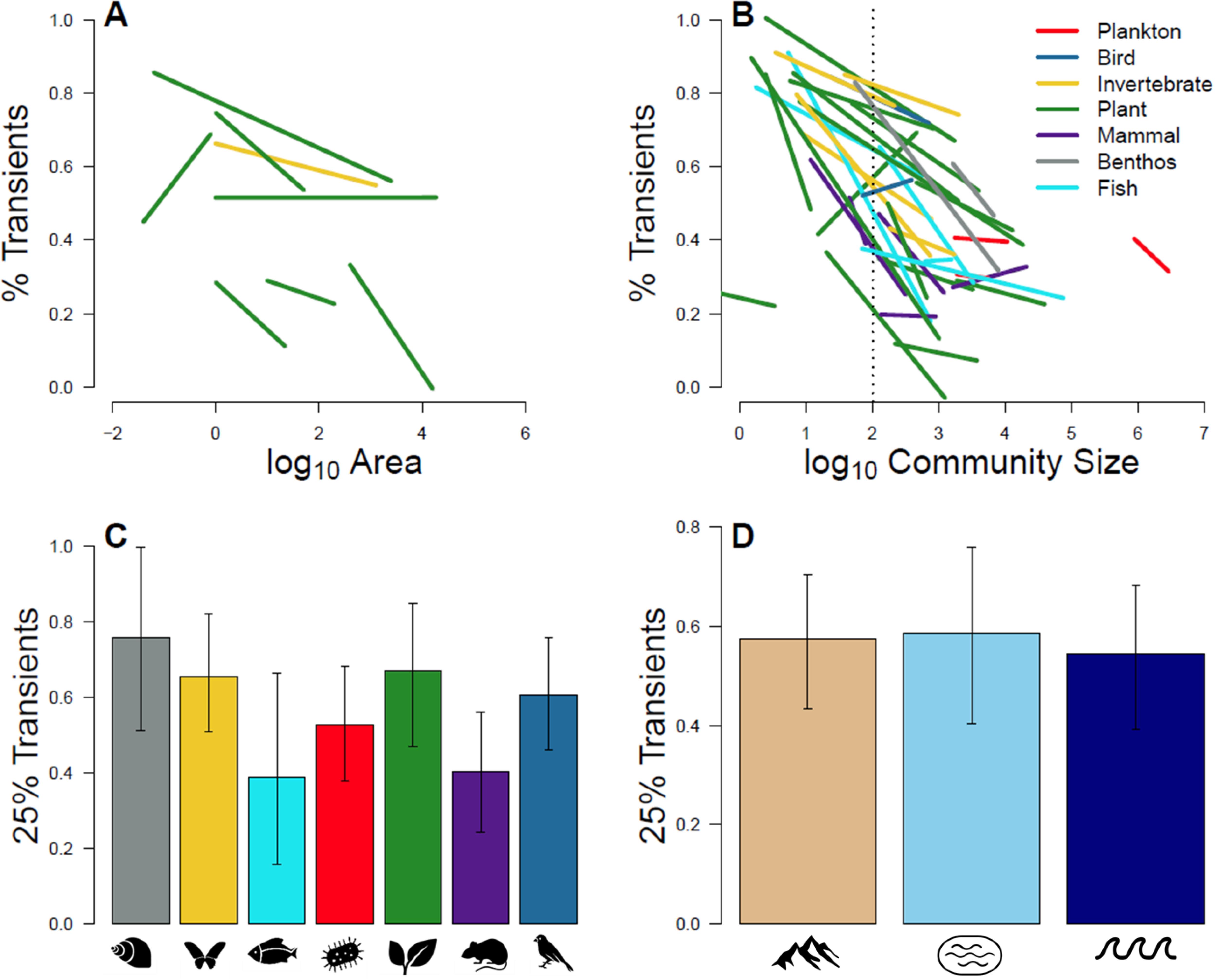
The impact of scale on the proportion of transient species as displayed in Figure 4 in the main text, but where transient species are defined as those with temporal occupancy ≤ 25%.

**Figure A5.**
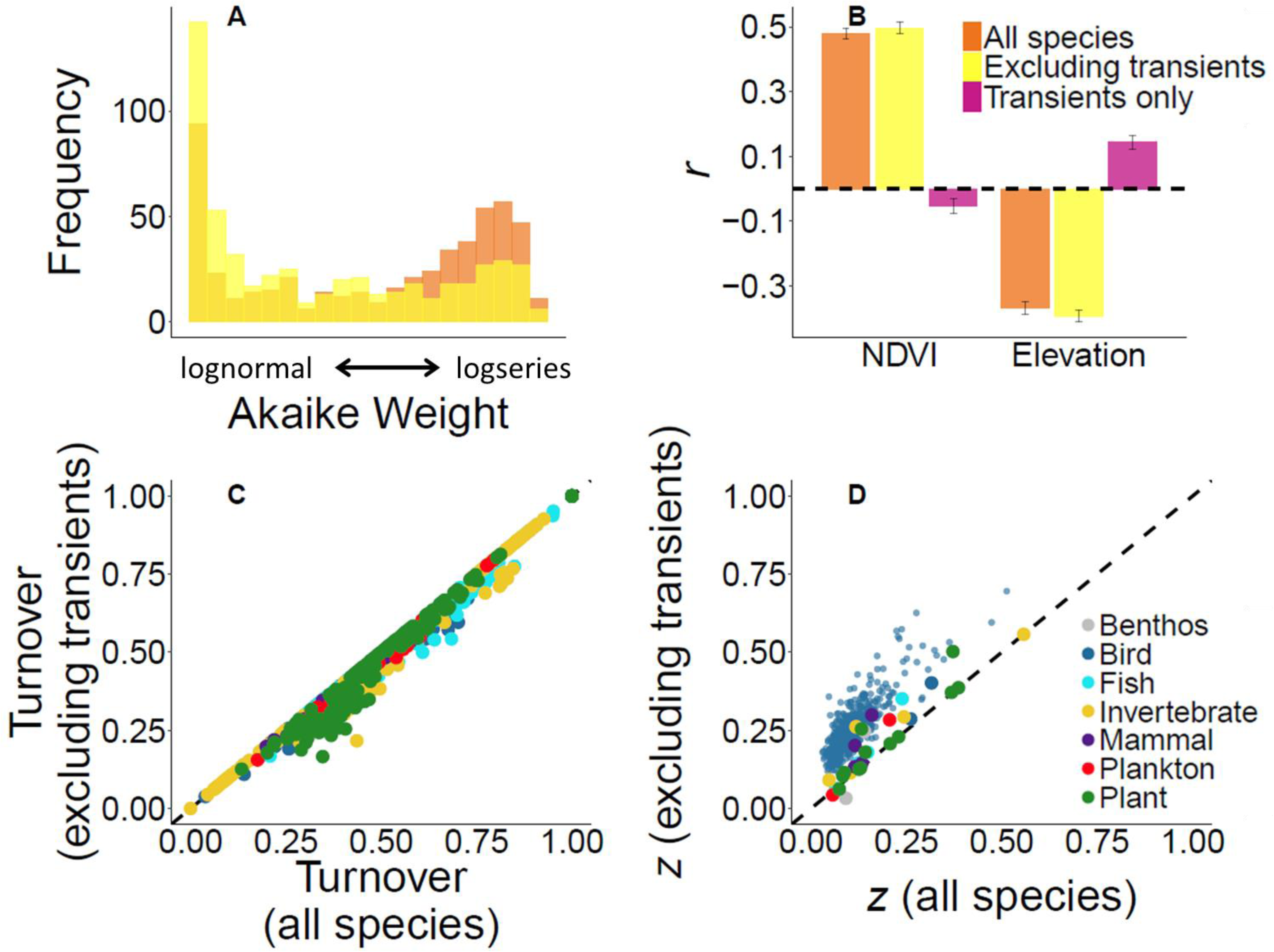
Impact of excluding transient species on four ecological patterns as displayed in Figure 5 in the main text, but where transient species are defined as those with temporal occupancy ≤ 10%.

**Figure A6.**
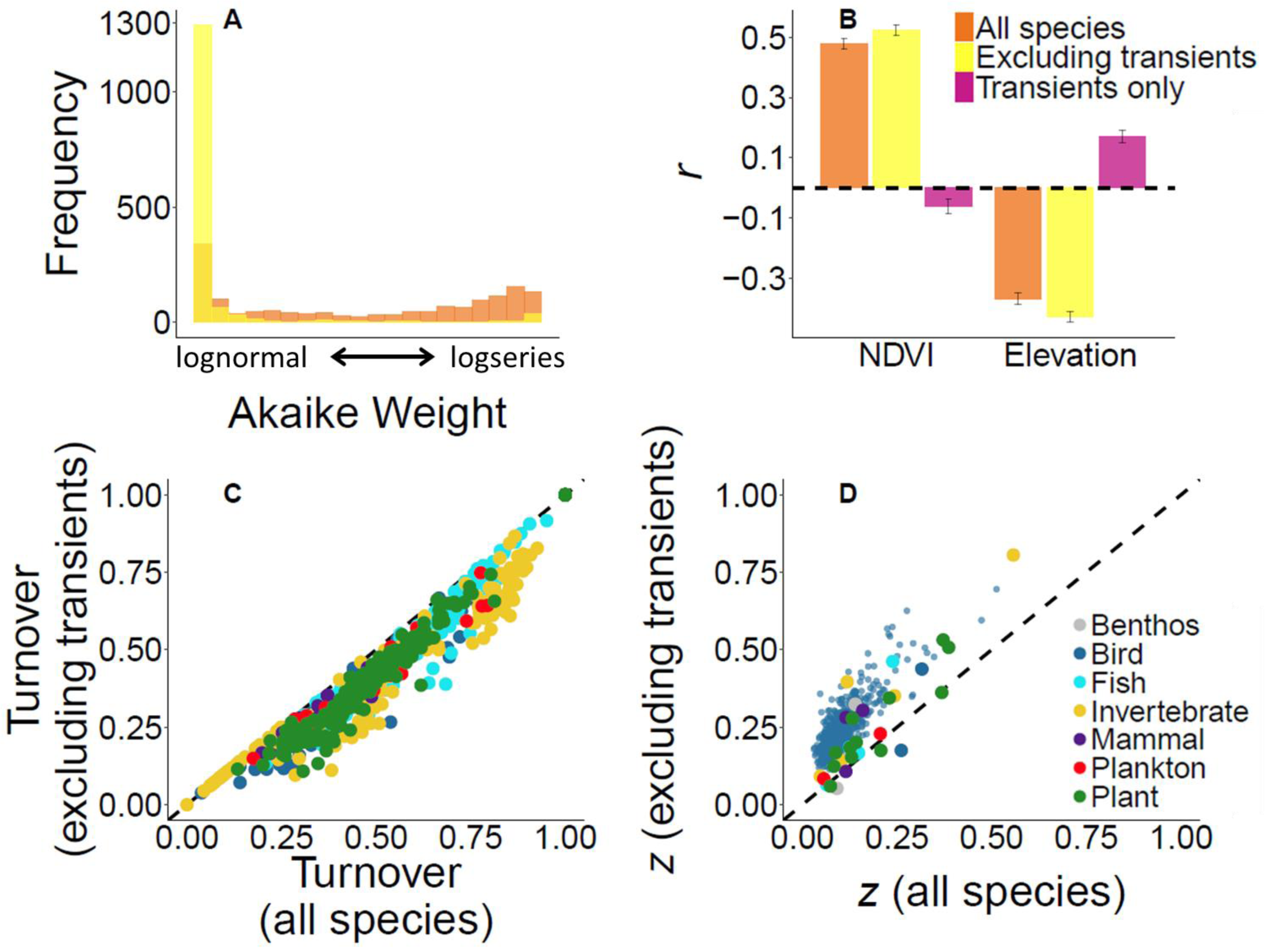
Impact of excluding transient species on four ecological patterns as displayed in Figure 5 in the main text, but where transient species are defined as those with temporal occupancy ≤ 25%

